# The neuronal calcium sensor NCS-1 regulates the phosphorylation state and activity of the Gα chaperone and GEF Ric-8A

**DOI:** 10.1101/2022.12.09.519724

**Authors:** D Muñoz-Reyes, LJ McClelland, S Arroyo-Urea, S Sánchez-Yepes, J Sabín, S Pérez-Suárez, M Menéndez, A Mansilla, J García-Nafría, SR Sprang, MJ Sánchez-Barrena

## Abstract

The Neuronal Calcium Sensor 1 and Ric-8A coregulate synapse number and probability of neurotransmitter release. Recently, the structures of Ric-8A bound to Gα have revealed how Ric-8A phosphorylation promotes Gα recognition and activity as a chaperone and guanine nucleotide exchange factor. However, the molecular mechanism by which NCS-1 regulates Ric-8A activity and its interaction with Gα subunits is not well understood. Given the interest in the NCS-1/Ric-8A complex as a therapeutic target in nervous system disorders, it is necessary to shed light on this molecular mechanism of action at atomic level. We have reconstituted NCS-1/Ric-8A complexes to conduct a multimodal approach and determine the sequence of Ca^2+^ signals and phosphorylation events that promote the interaction of Ric-8A with Gα. Our data show that the binding of NCS-1 and Gα to Ric-8A are mutually exclusive. Importantly, NCS-1 induces a profound structural rearrangement in Ric-8A that traps the protein in a conformational state that is inaccessible to Casein Kinase II-mediated phosphorylation, demonstrating one aspect of its negative regulation of Ric-8A-mediated G-protein signaling. Functional experiments indicate a loss of Ric-8A GEF activity towards Gα when complexed with NCS-1, and restoration of nucleotide exchange activity upon increasing Ca^2+^ concentration. Finally, the high-resolution crystallographic data reported here that define the NCS-1/Ric-8A interface will allow the development of therapeutic synapse function regulators with improved activity and selectivity.

## Introduction

Ca^2+^ is a key signal that regulates multiple biological phenomena ranging from neurotransmission to gene expression. Changes in the concentration of intracellular free Ca ^2+^, its locus of action and the amplitude and duration of Ca^2+^ influx are essential to transmit information through the nervous system. The mechanisms by which these changes can bring about such diverse neural responses rely on the ability of Ca^2+^ sensors to decode Ca^2+^ signals (*1, 2*). The Neuronal Calcium Sensor (NCS) family of proteins has evolved to participate in specialized neuronal functions separate from calmodulin, due to their 10-fold higher affinity for Ca^2+^. The most abundant protein of the NCS family is the Neuronal Calcium Sensor 1 (NCS-1), which was first discovered in drosophila and named Frequenin (Frq), is N-terminally myristoylated (Myr) and highly conserved from yeast to humans (*2–5*).

Unlike other NCSs, NCS-1 is found outside the nervous system. It does not contain a Ca^2+^/Myr switch, thus being constantly bound to the membrane, and has multiple binding partners (*2, 4– 6*). NCS-1 participates in a wide range of important neuronal functions: it is a regulator of Ca^2+^ channels, exocytosis, synaptogenesis and axonal growth, affecting higher functions such as learning and memory, neuroprotection and axonal regeneration (*2, 5, 7–10*). Furthermore, NCS-1 has been implicated in several pathological processes such as X-linked mental retardation and autism, schizophrenia and bipolar disorder (*11–14*). The multifunctionality of NCS-1 relies on its ability to recognize and regulate different and unrelated target proteins: G protein-coupled receptors (GPCRs) and some of their regulators, Ca^2+^ channels, guanine nucleotide exchange factors, and kinases, both in a Ca^2+^ dependent or independent manner (*2*).

The structure of NCS-1 consists of 2 pairs of EF-hand motifs, of which only three are functional: EF-2, EF-3 and EF-4 (*14*). EF-2 and EF-3 can recognize Mg^2+^ as well. It is known that the two Ca^2+^/Mg^2+^ binding sites are structural sites that allow the protein to adopt its tertiary fold. EF-4 has been suggested to be a regulatory Ca^2+^ binding site, able to sense changes in cytosolic calcium levels in neurons (*2, 15–20*).

The structures of several NCS proteins bound to their corresponding targets have shown that these Ca^2+^ sensors use a surface-exposed hydrophobic crevice to recognize their targets, which generally present small helical motifs that bind to the N- or C-terminal part of this large cavity (Fig 1A). It has been proposed that the structural determinants of target specificity are based on the shape and size of the hydrophobic crevice. NCS proteins contain a dynamic C-terminal helix (the so-called helix H10) that can insert into the crevice, thus contributing to its shape (Fig 1A). Since Ca^2+^ binding promotes structural rearrangements (Fig 1A) the occupancy of the three Ca^2+^ binding sites also determines affinity for protein partners. Also, the presence of hydrophilic residues at the border of the crevice contribute to target specificity and constitute hot spots for interactions with the different targets (*2, 6*).

**Figure 1:**
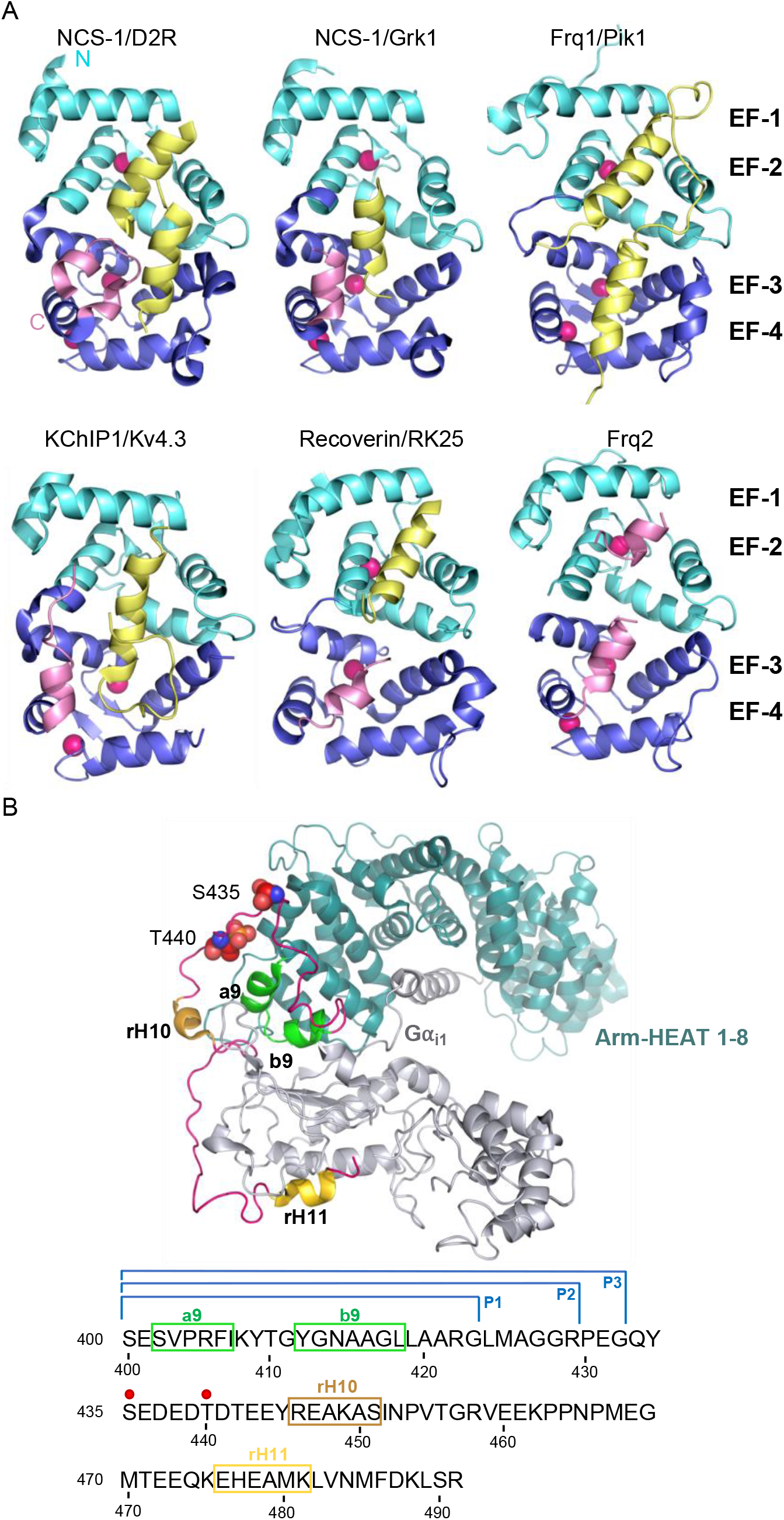
The structure of NCS/target complexes. **A)** Ribbon representation of NCS protein structures bound to their targets. NCS-1/DR2 (PDB: 5AER (*23*)), NCS-1/Grk1 (PDB: 5AFP (*23*)), Frq1/Pik1 (PDB: 2JU0 (*42*)), KChIP1/Kv4.3 (PDB: 2I2R (*54*)), Recoverin/RK25 (PDB: 2I94, (*78*)), Frq2 (PDB: 4BY4, (*25*)). The N and C-terminal pairs of EF-hands (EF-1-2 and 3-4) are shown in cyan and purple respectively. The C-terminal helix H10 is shown in pink and target proteins in yellow. Ca^2+^ is shown as hot pink spheres. **B)** Top: Cryo-EM structure of the rRic-8A/Gα_i1_ complex (PDB: 6UKT, (*39*)). Gα_i1_ is depicted in silver. Ric-8A Arm-HEAT repeats 1-8 in blue and repeat 9 in green and helices H10 (rH10) and H11 (rH11) in orange and gold respectively. The C-terminal coiled regions are shown in magenta. Phosphorylated residues S435 and T440 are depicted as spheres. Bottom: rRic-8A sequence from residue 400 to the end. Helices are squared following the same colour code used above. Phosphorylation sites are indicated as red spheres. P1, P2 and P3 brackets indicate the boundaries of the synthesized Ric-8A peptides.

Interestingly, NCS-1 is an important regulator of G protein signaling since it binds to proteins such as GPCRs including the Dopamine D2, Adenosine 2A and Cannabinoid CB1 receptors (*21–24*); and the molecular chaperone and Guanine nucleotide Exchange Factor (GEF) Ric-8A (*25*). Although little is known of the regulatory activity of NCS-1 on GPCR function and the cellular and physiological consequences, Romero-Pozuelo et al (2014) (*25*) showed that the NCS-1/Ric-8A complex is implicated in the regulation of synapse number and probability of neurotransmitter release. In fact, the regulation of this protein-protein interaction (PPI) using small molecule modulators allows synapse function control under pathological conditions. In neurodevelopmental disorders, where synapse number is abnormally high, the inhibition of the NCS-1/Ric-8A complex formation reduces synapse number and improves learning in Fragile X syndrome animal models (*26*). In contrast, the stabilization of the PPI prevents synapse loss and the consequent impairment in locomotion in a drosophila model of Alzheimer’s disease neurodegeneration at the motor neurons (*27*).

Ric-8A is an ubiquitously expressed cytosolic protein with two main functions: it constitutes a molecular chaperone that allows heterotrimeric Gα subunit biogenesis (*28*) and additionally, works as a guanine exchange factor for G_i,_, G_q_ and G_12/13_ families (*29–31*). Both activities are stimulated by Casein Kinase II (CK2) phosphorylation (*32*). Ric-8A has been shown to regulate asymmetric cell division and is essential for embryonic development (*33–37*). Work in drosophila has shown the relevance of Ric-8A in activating G_s_ for *in vivo* synaptogenesis and that this activity is downregulated by NCS-1 (*25*). Recently, the structure of several Ric-8A/Gα or Gα fragment complexes have been solved at atomic level (*38–40*). These works revealed the structural basis of Ric-8A as a Gα chaperone and GEF and showed how phosphorylation of Ric-8A residues S335 and T440 stabilize a conformation that is competent for Gα recognition. However, there is scarce information on the molecular function of NCS-1 on Ric-8A activity. Based on genetic studies, it has been proposed that NCS-1 interacts with Ric-8A and prevents the Ric-8A/Gα interaction (*25*). Here, we have combined biochemical, biophysical, crystallographic and cryo-EM studies to reveal the structural determinants of NCS-1/Ric-8A recognition and the mechanism of NCS-1-mediated downregulation of Ric-8A activity. This work shows how NCS-1 and Ric-8A constitute a hub that integrates Ca^2+^, phosphorylation and G-protein signaling. The emergent picture indicates that Ric-8A activity is under NCS-1 control and that a Ca^2+^ signal triggers the disassembly of the NCS-1/Ric-8A complex, which in turn allows phosphorylation of Ric-8A, which stabilizes the Ric-8A/Gα complex.

## Results

### The NCS-1 interacting region of Ric-8A and the role of Ca^2+^

To identify potential NCS-1 binding regions, we exploited the high-resolution structural information available on Ric-8A and NCS-1. An analysis of the different reported Ric-8A structures was performed to find potential NCS-1 binding regions (*39, 41*). First, we took into account that NCS protein targets generally employ one or two small helical motifs to recognize the N- or C-terminal pair of EF-hands (Fig 1A) (*2*). Second, we considered that the potential interacting helix or helices may have hydrophobic character, since Ric-8A interacts with NCS-1 through its surface-exposed hydrophobic crevice (*6, 25*). Third, NCS-1 and G proteins compete for Ric-8A binding and thus, could share certain interaction surfaces (Fig 1B) (*25*). Using these criteria, we evaluated the hydrophobic character of the HEAT repeat 9 of the ARM/HEAT repeat domain, which is composed of a two-helix bundle (called a9 and b9), as well as two C-terminal helical motifs (rH10 and rH11), all of them involved in Gα recognition (Fig 1B). The marked hydrophobic character of the a9-b9 two-helix bundle, led us to hypothesize that a9 and/or b9 helices could be implicated in the interaction with NCS-1.

To test our hypothesis, we carried out the *in vitro* reconstitution of the protein complex using an unphosphorylated Ric-8A rat variant construct ending at residue 452 (rRic-8A-452), which includes the two-helix bundle a9-b9 and rH10 (Fig 1B), and a human NCS-1 deletion construct (NCS-1ΔH10; 100% sequence identity to rat). The NCS-1 C-terminal helix H10 was removed since it works as a built-in competitive inhibitor of the Ric-8A interaction and the affinity for Ric-8A increases 30% in the absence of the helix (*25*). Previous co-immunoprecipitation assays with the NCS-1 drosophila variant suggested that the interaction occurred in the presence of CaCl_2_, although it was stronger in the presence of EDTA (*25*). To gain further insights into the Ca^2+^ dependency of the complex formation, reconstitution experiments were conducted at different Ca^2+^ concentrations and size exclusion chromatography (SEC) was used to evaluate complex assembly (see Materials and Methods section). Assembly (i) was attempted in Ca^2+^ free conditions (2mM EGTA with or without 1mM Mg^2+^; see Materials and Methods section). No sign of complex assembly was observed (Fig 2A). Assembly (ii) was carried out using a Ca^2+^ preloaded NCS-1 protein and maintaining 2 mM CaCl_2_ concentration in the SEC elution buffer. A very small peak at higher molecular weights was observed and most of the sample eluted as un-complexed species (Fig 2A). Therefore, neither a Ca^2+^ free nor a fully Ca^2+^ loaded NCS-1 protein recognized Ric-8A properly, suggesting that Ca^2+^ loading of some, out of the 3 Ca^2+^ binding sites, is required for recognition of Ric-8A. To achieve this, Ca^2+^-free EGTA-purified NCS-1 was mixed with Ric-8A at a final EGTA concentration of 0.6 mM, and subsequently dialyzed against a 2 mM CaCl_2_ buffer (Assembly (iii)). The majority of the protein sample eluted at an apparent molecular mass consistent with the formation of the NCS-1ΔH10/Ric-8A complex, as confirmed by SDS-PAGE gel (Fig 2A). Nano Differential Scanning Fluorimetry (nano-DSF) experiments also corroborated the efficient assembly when moving from EGTA to intermediate Ca^2+^ conditions (Fig 2B). In the presence of 0.6 mM EGTA, the denaturation profile of the mixture is consistent with the independent denaturation of both proteins, whose unbound forms have Ti values of 53.6 ºC (Ric-8A) and 59.5 ºC (NCS-1 EGTA) (Fig 2B). In contrast, the thermal transition of Ric-8A is up-shifted by more than 20 ºC (Ti = 77.5 ºC) in the sample dialyzed from EGTA to Ca^2+^ concentrations, due to the complex formation promoted by NCS-1 Ca^2+^ uptake. Moreover, the Ti of the Ca^2+^-bound form of NCS-1 in the sample is 85.5ºC, a value that is significantly lower than that shown when the three Ca^2+^ binding sites are saturated, which is above 90 ºC (Fig 2B).

**Figure 2:**
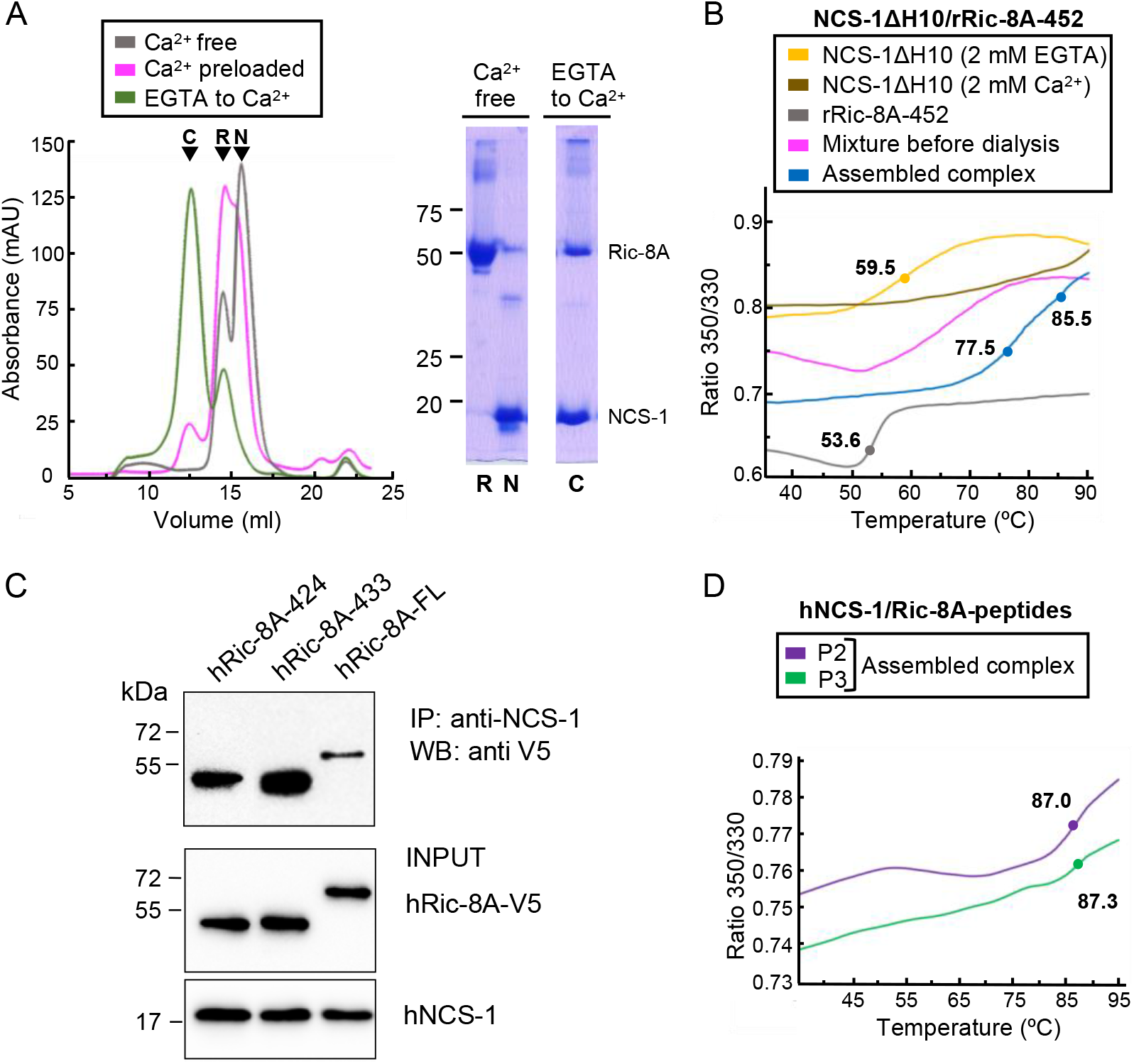
The assembly of rat and human NCS-1/Ric-8A complexes. **A)** Ca^2+^ dependency of the interaction of the rat complex. Size exclusion chromatograms after assemblies: (i) in Ca^2+^ free conditions (grey), (ii) with Ca^2+^ preloaded NCS-1ΔH10 (magenta) and (iii) with a dialysis from EGTA to Ca^2+^ (green). 12 % SDS-PAGE gels analyzing elution of NCS-1ΔH10 (N) and rRic-8A-452 (R) after assembly (i) and the NCS-1/rRic-8A-452 complex (C) after assembly (iii). **B)** Nano-DSF curves of the different samples during assembly (iii). The ratio between the emission fluorescence at 350 and 330 nm is shown *vs* the temperature. Curves corresponding to the EGTA purified NCS-1 (NCS-1 2 mM EGTA), the fully Ca^2+^ saturated protein (NCS-1 2 mM Ca^2+^) and rRic-8A-452 are shown as references in yellow, brown and grey, respectively. The mixture of proteins before dialysis (0.6 mM EGTA) and afterwards (assembled complex, 2 mM CaCl_2_) are shown in magenta and blue respectively. NCS-1 refers to NCS-1ΔH10, while Ric-8A to Ric-8A-452 construct. **C)** Co-IP protein-protein interaction assay in HEK293 cells of full-length human NCS-1 and V5-tagged hRic-8A constructs: full-length (hRic-8A-FL) and C-terminally truncated hRic-8A-424 (residue 1-424) and hRic-8A-433 (residue 1-433). **D)** Nano-DSF curves of hNCS-1 bound to different Ric-8A peptides. NCS-1 refers to NCS-1ΔH10, P2 and P3 refers to Ric-8A peptides P2 (purple) and P3 and (green).

The Ric-8A construct used in the *in vitro* reconstitution of the rat NCS-1/Ric-8A complex (rRic-8A-452) includes the two-helix bundle a9-b9 (HEAT repeat 9) and rH10 (Fig 1B). To determine whether elements beyond a9-b9 (e.g., rH10 and rH11) are implicated in NCS-1 recognition and also determine if the interaction occurs in the context of the human proteins, a co-immunoprecipitation assay was performed using both full-length human Ric-8A (hRic-8A-FL) and a C-terminally truncated construct lacking rH10 and rH11, and ending at G424 (hRic-8A-424, residues 1 to 424). (Fig 2C). The human Ric-8A sequence is one residue longer than the rat variant due to the insertion of a proline in a loop at position 208. Therefore, G424 in human corresponds to rat G423 (Fig 1B). Compared with hRic-8A-FL (Fig 2C), hRic-8A-424 has significantly higher affinity for NCS-1, suggesting that the HEAT repeat 9, but not the rH10 or rH11 helices, are implicated in the PPI. This supports a model in which two helices of Ric-8A are bound to NCS-1, similar to the case found for yeast NCS-1 bound to Pik1 (Fig 1A) (*42*). Also, this model is consistent with a reported crystal structure of drosophila NCS-1 (also known as Frq2) in its apo form (*25*), in which its hydrophobic crevice is occupied by two C-terminal NCS-1 H10 helices, one belonging to the same protein and an additional helix belonging to another molecule of the asymmetric unit, thus mimicking a protein/target complex (Fig 1A).

### The crystal structure of hNCS-1 bound to Ric-8A peptides

Attempts to crystallize the assembled *in vitro* reconstituted NCS-1ΔH10/rRic-8A complex were made with different rRic-8A constructs (ending at different positions between Ric-8A-423 and Ric-8A-452), without success. Therefore, we decided to work with peptides spanning the a9 and b9 helices (Fig 1B) to produce a minimal complex (named NCS-1/Ric-8A-P) and perform crystallographic studies. Three different peptides were synthesized starting at residue 400 and ending at position 423 (P1), 429 (P2) and 432 (P3) (Fig 1B). These peptides include a region of Ric-8A that is 100 % identical in both the human and rat variants. As indicated above, the human Ric-8A sequence is one residue longer than the rat variant due to the insertion of a proline in a loop at position 208. However, for easier comparison with the cryo-EM results presented in this work and previous structural studies carried out with the rat variant (*39, 41*) we have decided to maintain the rat numbering.

The assembly of the minimal complex was performed using conditions similar to those used to form assembly (iii) as described above. hNCS-1ΔH10 was incubated with the Ric-8A peptides in a 1:10 molar ratio and introducing Ca^2+^ by dialysis, starting at 1.7 mM EGTA and ending with a 0.5 mM Ca^2+^ concentration (see Materials and Methods). No crystals were obtained with the shortest peptide (P1). However, crystals were obtained with the complexes assembled with peptides P2 and P3. The crystals obtained with hNCS-1ΔH10/Ric-8A-P2 grew using microseeding techniques in conditions containing 0.5 mM Ca^2+^, 100 mM Mg^2+^ and 100 mM Na^+^ (see Materials and Methods section). hNCS-1ΔH10/Ric-8A-P3 crystals grew under similar conditions to those found for Ric-8A-P2. In addition, peptide P3 produced crystals in a second condition containing only Ca^2+^ and Na^+^ (see Materials and Methods section). Diffraction data sets were collected at the Spanish ALBA synchrotron (Table 1). All crystals belonged to the tetragonal space group P4_1_2_1_2 and displayed similar cell dimensions. The structure was solved by molecular replacement, using the Ric-8A-P2 data set (Structure 1, Table 1).

**Table 1:** Diffraction data collection and refinement statistics of hNCS-1/Ric-8A-P crystals.

| Data collection | Structure 1 | Structure 2 | Structure 3 |
| --- | --- | --- | --- |
| PDB code | 8ALH | 8AHY | 8ALM |
| Peptide | P2 | P3 | P3 |
| Ions in solution | Mg<sup>2+</sup>, Ca<sup>2+</sup>, Na<sup>+</sup> | Mg<sup>2+</sup>, Ca<sup>2+</sup>, Na<sup>+</sup> | Ca<sup>2+</sup>, Na<sup>+</sup> |
| Space group | P 4<sub>1</sub> 2<sub>1</sub> 2 | P 4<sub>1</sub> 2<sub>1</sub> 2 | P 4<sub>1</sub> 2<sub>1</sub> 2 |
| Cell dimensions |  |  |  |
| a, b, c (Å) | 56.86, 56.86, 134.61 | 56.64, 56.64, 135.30 | 56.64, 56.64, 134.53 |
| α, β, γ (°) | 90.00, 90.00, 90.00 | 90.00, 90.00, 90.00 | 90.00, 90.00, 90.00 |
| Resolution range (Å) | 52.38-1.86 (1.93-1.86)<sup>a</sup><br>[a*, b*=1.846, c*=1.917] | 52.24-1.70 (1.79-1.70)<sup>a</sup><br>[a*, b*=1.681, c*=1.891] | 52.20-1.85 (1.94-1.85)<sup>a</sup><br>[a*, b*=1.854, c*=1.920] |
| R<sub>pim</sub> | 0.044 (0.803) | 0.036 (0.659) | 0.028 (0.616) |
| CC<sub>1/2</sub> | 0.998 (0.445) | 0.997 (0.551) | 0.999 (0.467) |
| I / σI | 16.8 (1.2) | 13.5 (1.4) | 15.2 (1.3) |
| Completeness |  |  |  |
| Spherical (%) | 92.7 (41) | 88.0 (34.1) | 91.2 (36.1) |
| Elipsoidal (%) | 94.9 (51.5) | 95.9 (65.4) | 94.0 (45.7) |
| Wilson B-factor | 31.12 | 26.90 | 37.80 |
| Multiplicity | 25.2 (26.5) | 25.5 (27.7) | 8.7 (9.8) |
| **Refinement** |  |  |  |
| Resolution (Å) | 52.38-1.86 | 52.24-1.70 | 52.20-1.85 |
| No. reflections | 18029 | 22008 | 17665 |
| R<sub>work</sub> / R<sub>free</sub> | 19.58/23.13<br>(26.31/28.49) | 18.64/20.76<br>(34.65/45.00) | 20.98/25.25<br>(36.89/46.21) |
| **Assymmetric unit content** |  |  |  |
| No. atoms | 3454 | 3403 | 3374 |
| Protein (residue range) | 171 | 171 | 171 |
| Peptide (residue range) | 28 | 28 | 28 |
| PEG/GOL | 3/1 | 2/1 | 2/2 |
| Ca<sup>2+</sup>/Cl<sup>-</sup>/Mg<sup>2+</sup>/Na<sup>+</sup> ions | 2/1/1/1 | 2/1/1/1 | 2/1/0/2 |
| Water molecules | 141 | 144 | 116 |
| B-factor (Å)<sup>2</sup> | 31.27 | 27.28 | 37.18 |
| R.m.s deviations Protein |  |  |  |
| Bond lengths (Å) | 0.44 | 0.36 | 0.50 |
| Bond angles (°) | 0.63 | 0.56 | 0.65 |
| R.m.s deviations Peptide |  |  |  |
| Bond lengths (Å) | 0.45 | 0.57 | 0.33 |
| Bond angles (°) | 0.70 | 0.63 | 0.61 |

<sup>a</sup> Values in parenthesis are for highest resolution shell

Crystals of Ric-8A-P2 and -P3 each contain one complex in the asymmetric unit and both the 2F_o_-F_c_ and the F_o_-F_c_ electron density maps clearly indicated the presence of electron density corresponding to two helical segments (R1 and R2) of the peptide that completely occupy the NCS-1 hydrophobic crevice and protrude from its surface (Fig S1A and 3A). The quality of the data allowed the unambiguous modeling of Ric-8A-P2 residues 402-429; no density was found for the N-terminal residues S400 and E401 and the density corresponding to residues S402 to R405 was very weak (Fig 1B and S1A). The structures solved with Ric-8A-P3 (Structures 2 and 3, Table 1) were virtually identical with the exception that Mg^2+^ ion is present only in Structures 1 and 2. Compared with Structure 2, the C_α_ RMSD for the protein and peptide in Structures 1 and 3 were 0.090 and 0.227, and 0.103 and 0.117 respectively. The greatest differences are found between Structure 1 and 2 at the N-terminus of the Ric-8A peptides (Fig S1B), which make few contacts with NCS-1, and for which the temperature factors are high (Fig S1C). Although Ric-8A-P3 contains three extra residues at its C-terminus (Fig 1B) no electron density was found for the C-terminal residues P430, E431 and G432 suggesting that they are disordered, exposed to the solvent and do not participate in protein-protein recognition (Fig S1A). In fact, the thermal stability of NCS-1 bound to Ric-8A-P2 or Ric-8A-P3 is very similar (Fig 2D). The following discussion focuses on Structure 2, which is determined at the highest resolution and statistical quality (Table 1).

The structure of Ric-8A-P bound to NCS-1ΔH10 can be described as a coiled region followed by a short helical motif (R1) that is connected to a long helix (R2) through a loop (Fig 3A). The two helical elements are interconnected through the loop by an H-bond and van der Waals contacts (triad 1: I407-T410-A415) (Fig 3A and B) and also with helix-helix van der Waals interactions (F406-L418 and triad 2: K408-Y409-N414) (Fig 3A and B). Polar contacts with a chloride ion appear to stabilize the conformation of the turn between the two helical segments (Fig 3A). A calculation of the surface electrostatic potential of Ric-8A-P shows that the helices are amphipathic and expose positive charges to the solvent, except for the C-terminal tip, which shows a negative potential due to the carboxylic end of the peptide (Fig 3C).

**Figure 3:**
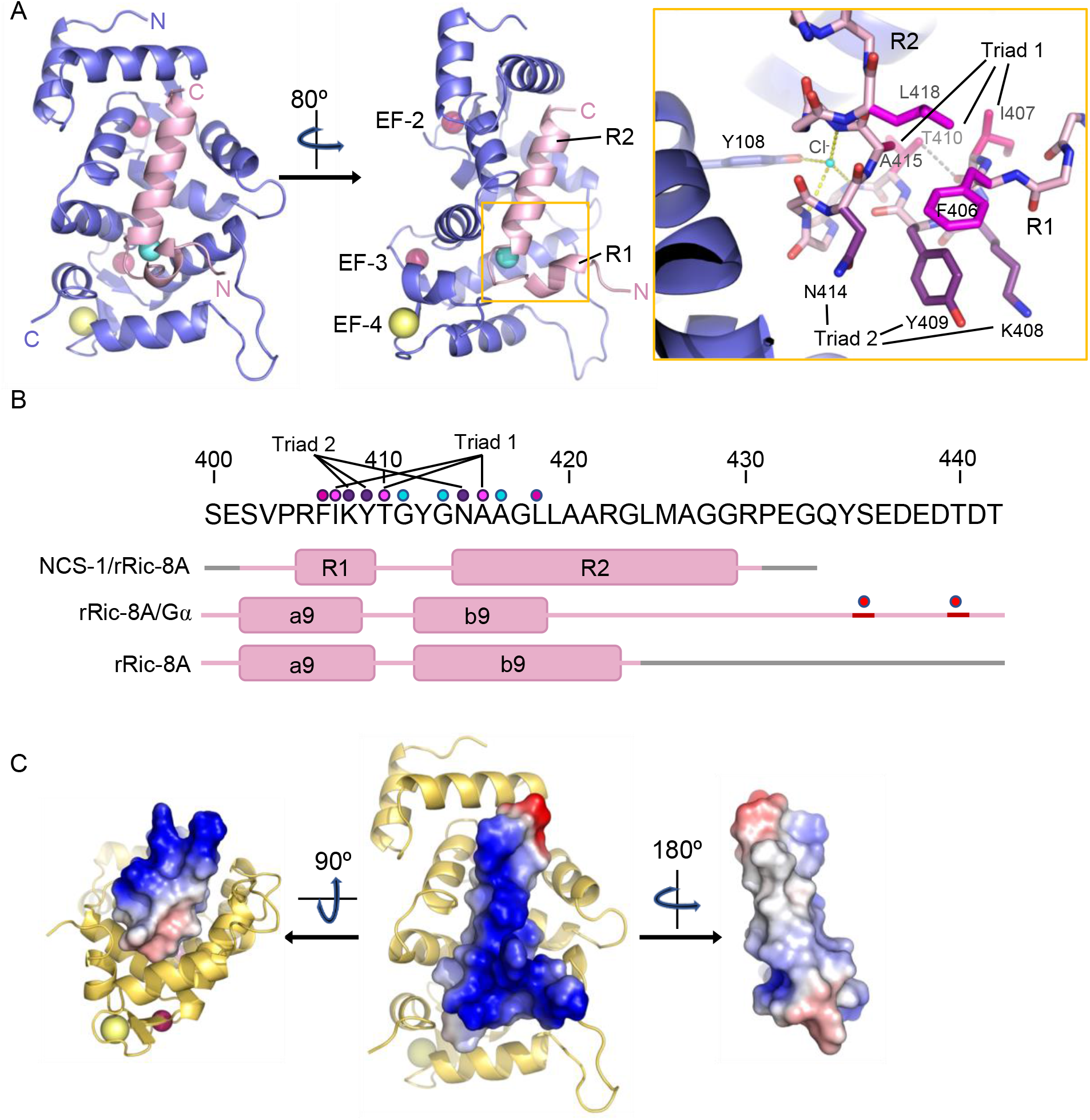
The structure of hNCS-1 bound to a Ric-8A peptide. **A)** Ribbon representation of the hNCS-1ΔH10/Ric-8A-P3 complex. Two views are displayed. The NCS-1 structure is shown in light purple, while Ric-8A-P3 is shown in light pink. The N- and C-termini are indicated. Ca^2+^, Na^+^ and Cl^-^ ions are shown in hotpink, yellow and cyan, respectively. R1 and R2 helices, and EF-hands 2, 3 and 4 are indicated. The orange square represents a zoomed view of the R1-R2 loop in stick mode, Cl^-^ coordination and H-bonds are displayed as yellow and grey dashes, respectively. Residues participating in R1-R2 contacts are displayed in hot pink (Triad 1: I407-T410-A415), magenta (F406-L418) and purple (Triad 2: K408-Y409-N414). **B)** rRic-8A sequence from 400 to 442 residues. The helix boundaries of Ric-8A sequence encompassing a9 and b9 in different structural contexts (NCS-1/Ric-8A-peptide (PDB: 8AHY), Ric-8A/Gα (PDB: 6UKT, (*39*)) and uncomplexed Ric-8A (PDB: 6NMG, (*41*)) are indicated as pink boxes and labelled. Coiled regions are shown in pink. Disordered regions are shown in grey, while phosphorylated sites are shown as red spheres. The interacting residues shown in panel (A) are indicated with dots in the same color code as in (A). **C)** Electrostatic surface potential of rRic-8A-P3. NCS-1 is shown as yellow ribbons. Positive and negative potential are represented in blue and red. On the right, the Ric-8A region that faces and contacts NCS-1 is shown with NCS-1 removed for proper visualization.

The hNCS-1ΔH10/Ric-8A-P contact area is 1140 Å^2^ (*43*) and a total of 1469 contacts take place (4.2 Å cut-off distance). All the NCS-1 helices that shape the cavity are implicated in Ric-8A recognition: while R2 helix is recognized by multiple helices, R1 contacts helices H7 and H8 (Fig S2A and S2B). A total of nine H-bonds are formed with Ric-8A in the cavity (Fig S2B). Helix R2 is recognized at the middle (L419), N-(N414) and C-terminus (R429) through H-bonds. The Ric-8A R1-R2 loop also plays a relevant role in the recognition process and Y412 forms two water-mediated H-bonds with residues located at the C-terminal part of the crevice (Fig S2B and S2C). NCS-1 residue R148 uses its guanidinium group to establish two direct H-bonds with T410 (R1-R2 loop) and K408 (R2 helix). All NCS-1 residues involved in polar contacts are located at the border of the hydrophobic crevice in which Ric-8A is inserted (Fig S2B). In addition to the polar contacts, a large number of van der Waals interactions are formed with the bottom and lateral walls of the NCS-1 crevice in which hydrophobic and a good number of aromatic residues are present (Fig S2A). Several hydrophobic and aromatic Ric-8A residues satisfy these interactions (Fig S2D).

Using Ric-8A peptides of different lengths to generate crystal structures, together with protein-protein interaction assays, it has been possible to define the region of Ric-8A that is necessary and sufficient for recognition of NCS-1 and which is conserved in rat and human protein sequences. The addition of three extra residues (peptide P3 vs P2), may allow better folding of the R2 helix. In fact, a cell-based protein-protein interaction assay with an hRic-8A deletion mutant (hRic-8A-433) ending like rRic-8A-P3 at G432 (Fig 1C) shows the strongest interaction with full-length hNCS-1 (Fig 2C). The crystal structure shows that all Ric-8A-P modelled residues except the N-terminal S402, and the residues P404, R405 and F406 (which belong to helix R1 and are exposed to the solvent, opposite to the face that contacts NCS-1) (Fig S1A, 3B, S2B-D) are involved in the interaction. Although helix R2 highly contributes to the protein-protein contact area, the R1-R2 loop, formed by residues T410, G411, Y412 and G413, plays an important role as well, contributing three hydrogen bonds (Fig S2). In fact, these four residues account for 28 % of total contacts between hNCS-1 and Ric-8A-P.

### The Ca^2+^ binding sites of hNCS-1 in complex with Ric-8A-P

The analysis of the structure of the hNCS-1ΔH10/Ric-8A-P complexes presented in this work, together with electron density map calculations, indicate that the three Ca^2+^ binding sites, EF-2, EF-3 and EF-4, are occupied, showing a pentagonal-bipyramidal coordination (Fig 4A). However, the *in vitro* reconstitution assays presented above (Fig 2A) show that the fully Ca^2+^ saturated protein does not efficiently generate the NCS-1/Ric-8A complex. Therefore, some out of the 3 sites are not occupied with Ca^2+^. NCS-1 Ca^2+^ occupancy has been addressed previously with the calculation of difference anomalous maps since Ca^2+^ but not Mg^2+^ or Na^+^ shows anomalous signal at typical protein-diffraction wavelengths (*6*). In fact, at 0.979 Å wavelength, the anomalous scattering coefficients (f’’) for Ca^2+^, Na^+^ and Mg^2+^ are 0.616, 0.049 and 0.073 electrons, respectively. Therefore, to identify the Ca^2+^ ions bound to the EF-hands, anomalous difference maps were calculated as a 10-fold increased anomalous signal is expected if Ca^2+^ is present. We used the best quality data set (Structure 2), in which Ca^2+^, Na^+^ and Mg^2+^ where present in the crystallization solution. The anomalous difference map shows 6 σ peaks at EF-2 and EF-3 metal sites while no signal is observed in EF-4, indicating unambiguously the presence of Ca^2+^ at sites EF-2 and EF-3 (Fig 4A). EF-4 shows a residual anomalous signal when reducing the σ level, which could indicate a low occupancy of Ca^2+^ at EF-4, consistent with the observation that a fully saturated NCS-1 does not recognize Ric-8A properly (Fig 2A). We have not modelled Mg^2+^ at any of the metal binding sites since it has been reported that EF-4 is unable to bind this metal (*15*). Furthermore, the metal-oxygen distances observed (2.3-2.4 Å) are higher than those characteristic of Mg^2+^ coordination (2.1 Å) (*44, 45*). We did observe a Mg^2+^ ion at the surface of the NCS-1, close to EF-3 (Fig 4A) and located at a special position where atoms from two symmetry related molecules participate in the coordination sphere. In this case, Mg^2+^ exhibits typical octahedral coordination and metal-oxygen distances (2.1 Å), as well as the absence of an anomalous signal. Taking these observations together, a Na^+^ ion was modelled at EF-4 (Fig 4A), since Na^+^ was present in the crystallization solution, does not scatter X-rays anomalously and is structurally undistinguishable from Ca^2+^. Their similar ionic radii allow equivalent heptahedral geometry, distances and angles (*45, 46*).

**Figure 4:**
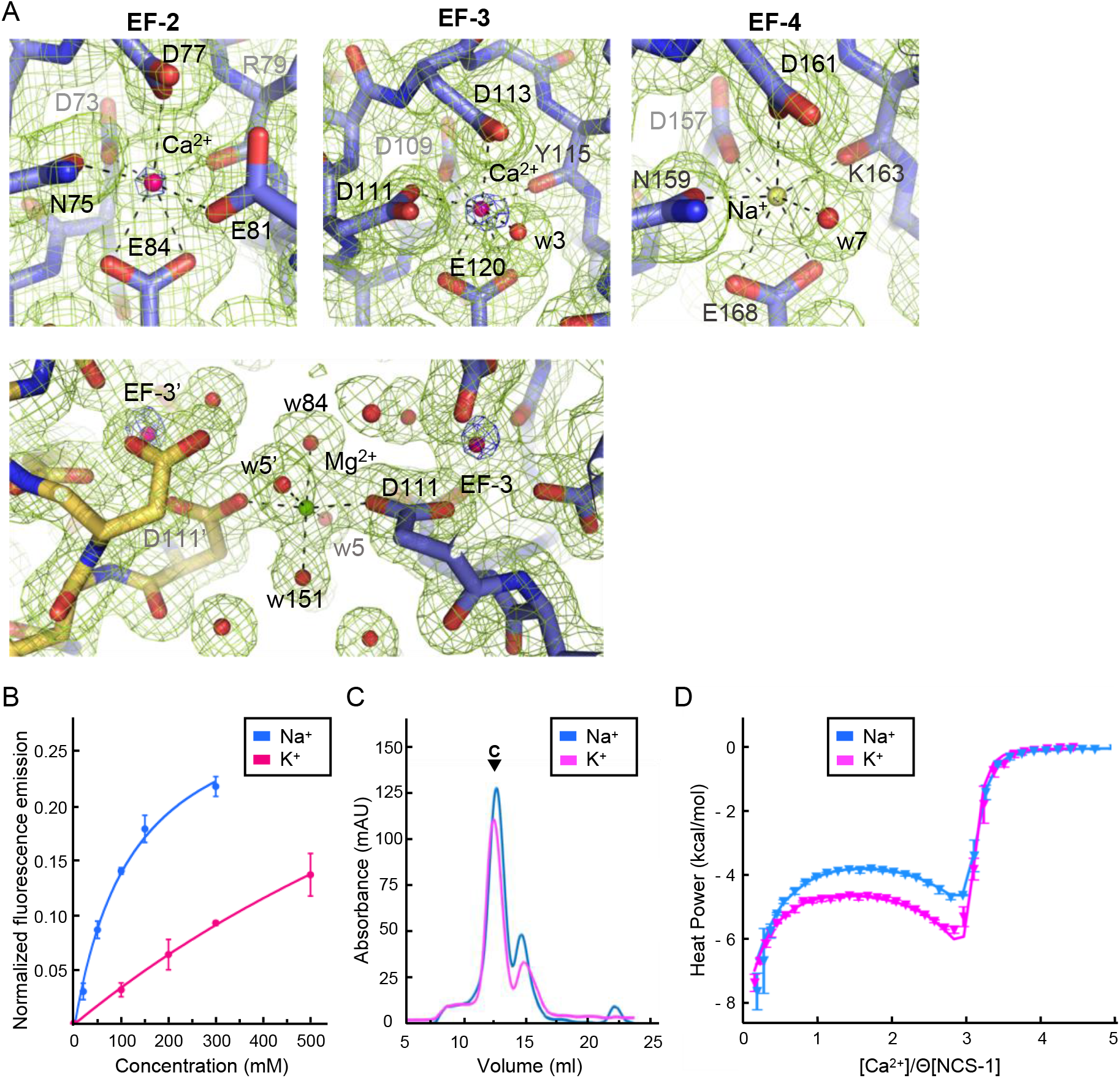
NCS-1 Ca^2+^ binding sites. **A)** Identification of Ca^2+^, Mg^2+^ and Na^+^ ions in the hNCS-1ΔH10/Ric-8A-P3 complex (Structure 2, see Table 1). Top: Electron density at EF-hands EF-2, 3 and 4. The 2F_o_-F_c_ electron density map (contoured at 1.0 σ) and the anomalous difference map (contoured at 6.0 σ) are shown in green and blue respectively. NCS-1 is shown in stick mode (light purple), Ca^2+^ and Na^+^ ions as hot-pink and yellow spheres, respectively, and water molecules (w) as red spheres. Bottom: The Mg^2+^ ion (green sphere) found in Structure 1 and 2 (see Table 1). NCS-1 symmetry related molecule is depicted in yellow. **B)** The binding of Na^+^ to hNCS-1 in solution. Representation of the normalized fluorescence emission (mean ± SEM; *n*=3) of full-length hNCS-1 at increasing concentrations of NaCl or KCl. The curves are the least squares fitting of the experimental data to a 1:1 stoichiometry equilibrium. Na^+^ and K^+^ titrations are shown in blue and magenta respectively. **C)** Assembly of the NCS-1ΔH10/rRic-8A-452 complex in the presence of 200 mM Na^+^ (blue) or K^+^ (magenta). Size exclusion chromatograms indicating the elution of the assembled complexes (C). **D)** ITC binding isotherm at 25 ºC for Ca^2+^ to NCS-1 in 20 mM Tris pH 7.9 supplemented with 150 mM Na^+^ (blue) or 150 mM K^+^ (magenta). Solid lines show the best fits of the titration data in terms of a three-site sequential binding model using the thermodynamic parameters shown in Table 2. Θ is the faction of sites available for each class of Ca^2+^ sites.

EF-hand containing proteins, for example, parvalbumin, are able to bind Na^+^ at their Ca^2+^ binding sites (*47*). Using tryptophan emission fluorescence experiments, we studied the ability of full-length hNCS-1 to bind Na^+^ in solution (Fig 4B). As a control, we also studied the binding to K^+^. Our data show that an increase in Na^+^ concentration produces an enhancement of the emission fluorescence intensity that achieves saturation at 300 mM NaCl. Considering a 1:1 equilibrium, it is possible to estimate the apparent dissociation constant from these data, which is 123.4 ± 26.6 mM (*6*). Compared with Na^+^, the addition of K^+^ promotes smaller changes in the emission intensity, which varied linearly with the cation concentration up to 500 mM KCl, with no sign of saturation. This suggests that NCS-1 is unable to bind K^+^ at physiologically relevant concentrations (Fig 4B). Therefore, the fluorescence experiments support the binding of Na^+^ to NCS-1.

**Table 2:** Thermodynamic parameters of Ca^2+^ binding to full-length hNCS-1 in the presence of K^+^ or Na^+^.

| **C(150mM)** | **$K_{d1}$ (nM)** | **$\Delta H_1$**<br>**(kcal/mol)** | **$K_{d2}$ (nM)** | **$\Delta H_2$**<br>**(kcal/mol)** | **$K_{d3}$ (nM)** | **$\Delta H_3$**<br>**(kcal/mol)** |
| --- | --- | --- | --- | --- | --- | --- |
| **Na<sup>+</sup>** | 265 ± 6 | -7.7 ± 0.1 | 758 ± 16 | 3.0 ± 0.1 | 379 ± 17 | -9.1 ± 0.3 |
| **K<sup>+</sup>** | 165.6 ± 0.3 | -7.66 ± 0.01 | 362.3 ± 0.6 | 1.00 ± 0,01 | 253 ± 1 | -9.44 ± 0,01 |

Subscripts 1, 2 and 3 correspond to sites 1, 2 and 3, respectively.

To determine whether the binding of Na^+^ to EF-4 is relevant for NCS-1/Ric-8A assembly, we compared the *in vitro* assembly of rat Ric-8A-452 and NCS-1ΔH10 in the presence of 200 mM NaCl or KCl and 2 mM Ca^2+^ (that is, the so-called assembly (iii) in the presence of NaCl or KCl). The formation of the complex is achieved in the presence of either ion (Fig 4C). Therefore, as suggested by the Ca^2+^ titration experiments (Fig 2A) only binding of Ca^2+^ to the structural sites EF-2 and EF-3 (*15*) triggers the conformational rearrangement needed for Ric-8A recognition and Na^+^ occupation of EF-4 does not contribute to NCS-1/Ric-8A complex formation. Therefore, Ric-8A can bind to NCS-1 species with a Ca^2+^-free EF-4 site, and in this situation, and at the 100 mM Na^+^ concentrations used in the crystallization conditions, the site is occupied with Na^+^.

Given that NCS-1 EF-4 binds Ca^2+^ (*14*) but also Na^+^, we used Isothermal Titration Calorimetry (ITC) to determine whether binding of Na^+^ to EF-4 affects the Ca^2+^-binding properties of NCS-1. We compared the binding isotherms obtained in the presence of 150 mM NaCl or KCl (control). In both media the binding data can be fit to a sequential binding model assuming three available sites (Fig 4D), with binding being enthalpy-driven for sites 1 and 3 and entropy-driven for site 2. As shown in Fig S3, filling of site 3 is significantly delayed compared to the occupation of the other two sites, which agrees with previous NMR-based data showing a sequential binding of Ca^2+^ to EF-2, EF-3 and finally EF-4 (*15–18*). It is also consistent with our crystal structure, where Ca^2+^ is found only in EF-2 and EF-3 sites (Fig 4A). A direct inspection of the isotherms suggests that the interaction with Ca^2+^ depends on the monovalent cation present in the media. In particular, we observe that the enthalpy of binding to site 2 (EF-3) triples in the presence of Na^+^, and the dissociation constants of sites 1 and 3 increase by about 1.5 times, while that of site 2 doubles. Therefore, our data suggest that Na^+^ binding modulates the affinity of NCS-1 for Ca^2+^, the influence likely extending to more than one site due to communication among them (*20*). However, the possibility that Na^+^ could bind to more than one unoccupied Ca^2+^ site cannot be completely discarded.

### The cryo-EM structure of NCS-1 bound to rRic-8A-452

Given the lack of success in the crystallization of the NCS-1ΔH10/rRic-8A-452 complex, we aimed to perform single-particle cryo-electron microscopy (cryo-EM). For this purpose we incubated the NCS-1ΔH10/rRic-8A-452 complex with nanobodies (Nbs) Nb8109, Nb8117 and Nb8119 raised against rRic-8A (*39*), which increased the molecular mass up to 115 kDa, making it amenable to cryo-EM analysis (see Materials and Methods). Cryo-EM grids were prepared with the freshly purified gel-filtered heteropentameric complex and data were acquired in a Titan Krios at the ESRF (see Materials and Methods section) (*48*). Upon data processing the complex displayed high levels of dissociation and conformational heterogeneity (see final 3D classification in Fig S4 for an example) limiting the resolution of the final map to 17 Å. However useful insights were obtained from these data.

The map showed a horseshoe shaped EM density into which the rRic-8A(1-400)/Nbs complex (PDB: 6UKT (*39*)) could be unambiguously fit using the Nbs as fiducial markers (Fig 5A, B and S5). The best docking position was also found for the hNCS-1ΔH10/Ric-8A-P crystal structure determined in this work within the remaining density (Fig 5A, inset) (see Materials and Methods and Fig S5B for docking analysis).

**Figure 5:**
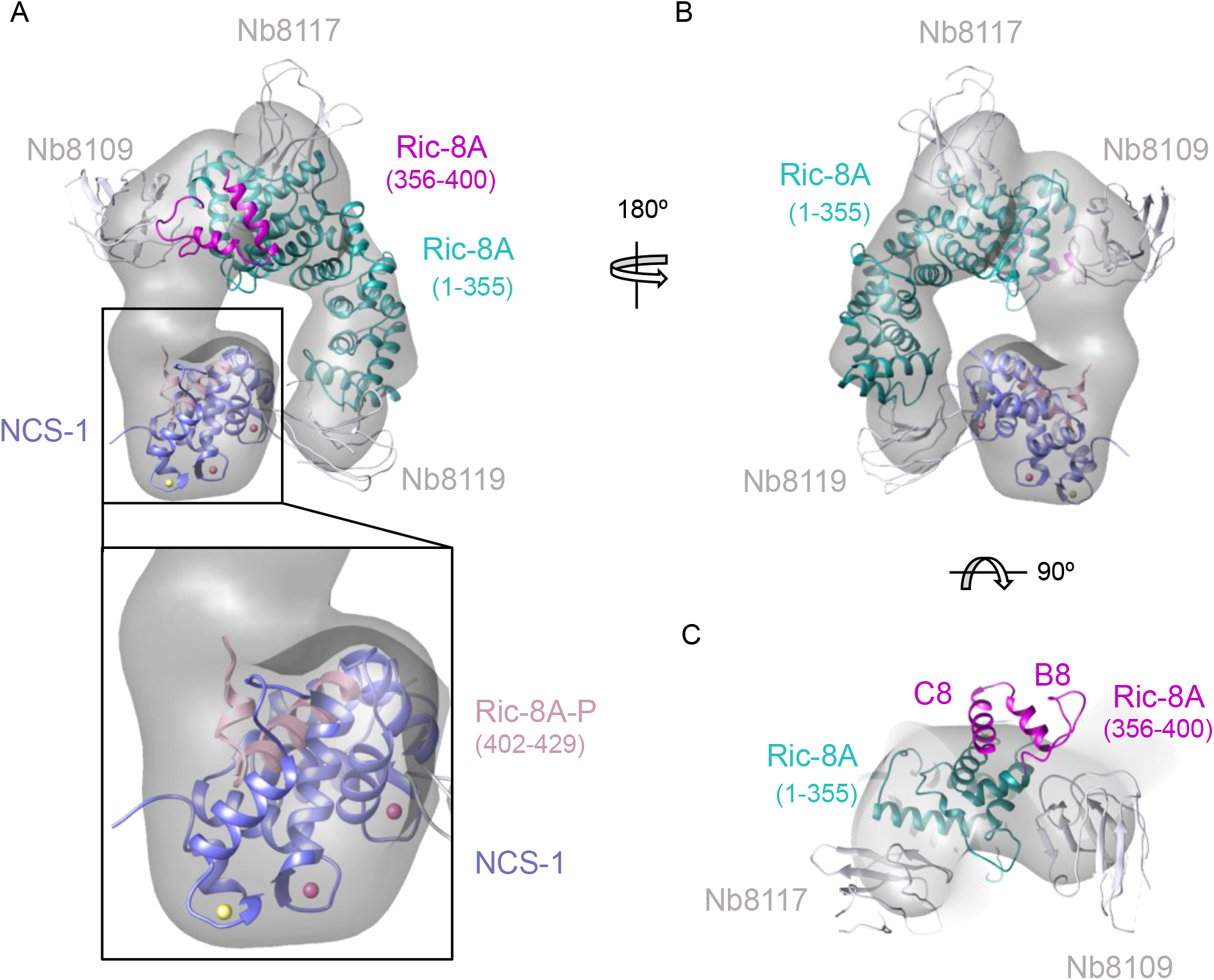
Cryo-EM map and fitting of rRic-8A and hNCS-1/Ric-8A-P coordinates. Front **(A)** and back **(B)** view of the cryo-EM map (grey) with the docked rRic-8A-400/Nbs (PDB: 6UKT, (*39*)) and hNCS-1/Ric-8A-P complexes (displayed as cartoons). The inset shows the N-terminus of Ric-8A-P protruding into the unassigned density. **C)** Top view of the complex showing a part of the HEAT repeat 8 (residue 356-400) protruding out of the cryo-EM map.

The final organization of the complex positions the C-terminal-most domain of rRic-8A(1-400) pointing towards the NCS-1ΔH10/Ric-8A-P complex, while the N-terminus of the Ric-8A-P in the NCS-1ΔH10/Ric-8A-P structure points towards the C-terminal region of rRic-8A, consistent with a connected complex. Between both fit coordinates there is an unaccounted volume of density in the cryo-EM map which displays poor signal at lower map thresholds, indicating that the connecting protein segment displays higher mobility in this region when compared to the rest of the protein. The arrangement of rRic-8A-452 and NCS-1 in the complex, together with the unaccounted density, suggest a reorganization of the C-terminal section of Ric-8A. Indeed, the ARM repeat 8, and more specifically, the long A8-B8 loop and helices B8 and C8 (residues 356 to 400), fall out of density in the cryo-EM map when docking the Ric-8A/Nbs complex as a whole (Fig 5C), which further supports the hypothesis of a conformational change at this part of Ric-8A. Furthermore, a crude fit of Ric-8A helices B8 and C8 into the unaccounted density shows a reasonable connection between both protein components in the complex, adopting an extended conformation (Fig S5A).

### Ric-8A phosphorylation in the context of the NCS-1/Ric-8A complex

Phosphorylation of rRic-8A at S435 and T440 residues promotes Gα subunits binding, since the interaction of the phosphorylated residues with a basic groove found at the ARM-HEAT repeat domain of Ric-8A creates a structural platform that allows Gα recognition (Fig 1B) (*38–40*). Here, we have shown that *in vitro*, the NCS-1ΔH10/rRic-8A complex assembly occurs with unphosphorylated rRic-8A. Thus, we have generated the corresponding non-phosphorylatable human full-length Ric-8A mutant (S436A, T441A; Ric-8A-P-Mut) and tested the binding to full-length hNCS-1 in a cell-based protein-protein interaction assay (Fig 6A). The co-immunoprecipitation of the protein complex shows that the non-phosphorylatable version of hRic-8A retains its ability to interact with hNCS-1. Interestingly, the binding is increased with respect to the wild-type version, which would suggest that Ric-8A is partly phosphorylated *in vivo* and this might hinder the interaction with NCS-1.

**Figure 6:**
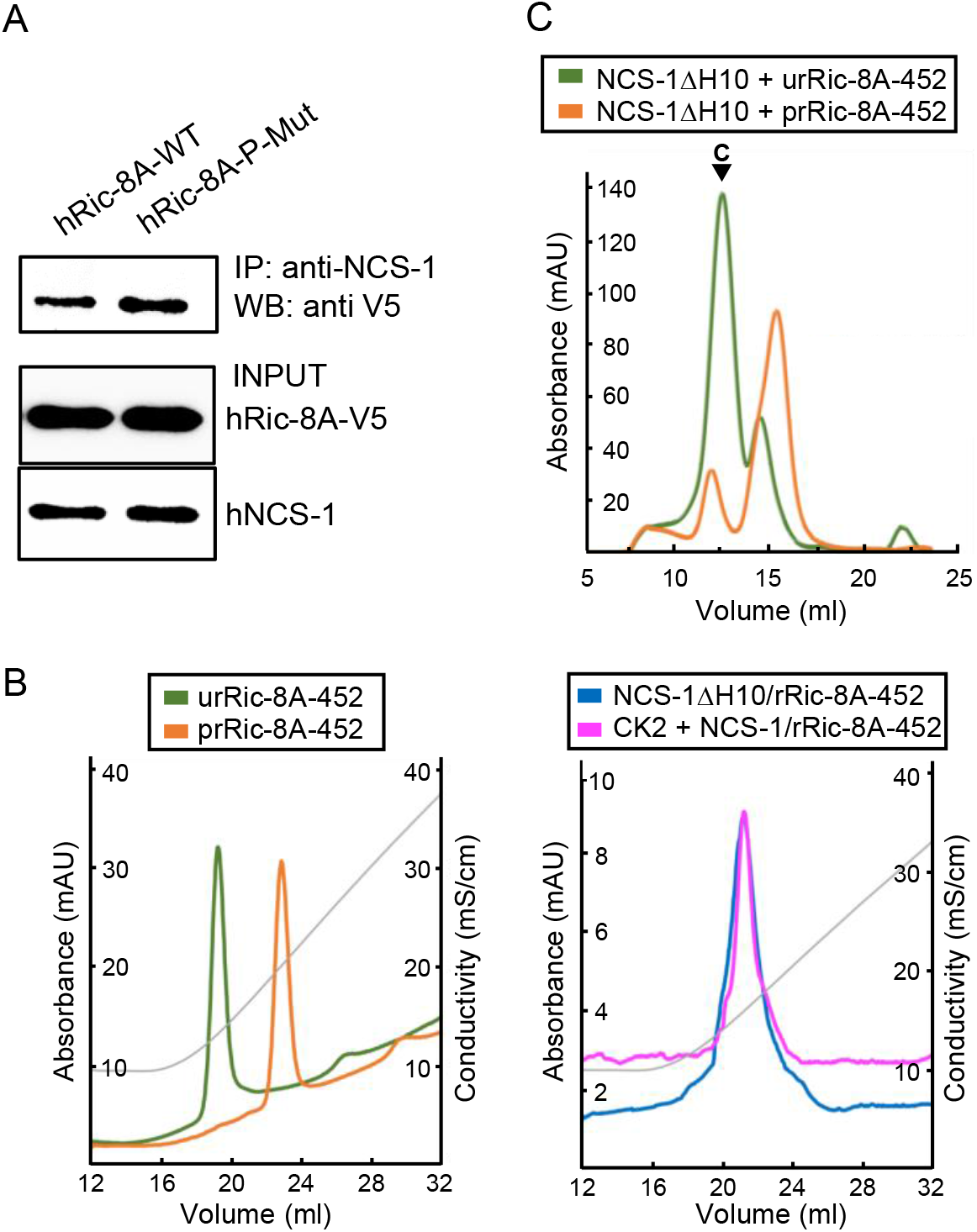
Ric-8A phosphorylation in the context of the NCS-1/Ric-8A complex. **A)** Co-IP protein-protein interaction assay of hNCS-1 and V5-tagged full-length hRic-8A WT (hRic-8A-WT) and a non-phosphorylatable mutant (Ric-8A-P-Mut; S436A, T441A) in HEK293 cells. **B)** Anionic exchange chromatograms of CK2 treated samples eluted in a salt gradient. On the left, phosphorylated and unphosphorylated rRic-8A-452 (prRic-8A-452 (orange) and urRic-8A-452 (green), respectively. On the right, CK2 treated (pink) or untreated (blue) NCS-1ΔH10/rRic-8A samples. Conductivity (mS/cm) is shown as grey lines. **C)** Size exclusion chromatograms of the resulting samples after the assembly of NCS-1ΔH10 with unphosphorylated (green) and phosphorylated (orange) rRic-8A-452. C stands for assembled complex.

To show whether NCS-1 has an influence on Ric-8A phosphorylation, we attempted to phosphorylate the NCS-1ΔH10/rRic-8A-452 complex using Casein Kinase II (CK2) following a protocol similar to that described by McClelland et al. (2020) (*39*). A control experiment was performed with the uncomplexed rRic-8A-452 protein. Protein phosphorylation was evaluated by anion exchange chromatography after incubation with CK2. The phosphorylated species elute at higher salt concentration since their anionic character is increased with the incorporation of the phosphate groups. As shown in Fig 6B, while uncomplexed rRic-8A-452 is fully phosphorylated after CK2 treatment, rRic-8A-452 is not phosphorylated when complexed with NCS-1, since the anionic exchange elution profile of the NCS-1ΔH10/rRic-8A-452 complex is the same regardless CK2 treatment. To further verify the phosphorylation state of Ric-8A in the CK2 treated NCS-1ΔH10/rRic-8A-452 complex, we used LC-MS/MS with the aim of identifying phosphopeptides from a trypsin-digested sample (see Material and Methods). No phosphopeptides were found even though a phosphopeptide enrichment protocol was performed, supporting the hypothesis that the binding of NCS-1 precludes CK2-mediated phosphorylation of Ric-8A.

Finally, we asked whether phosphorylated Ric-8A can interact with NCS-1. An assembly experiment was performed similarly to that described for the unphosphorylated protein (control). Our data show that phosphorylated rRic-8A-452 does not efficiently interact with NCS-1ΔH10 (Fig 6C).

### NCS-1/Ric-8A nucleotide exchange functional assays

To directly measure the effect of NCS-1 on Ric-8A GEF activity, nucleotide exchange assays were performed in the presence of increasing concentrations of CaCl_2_. With the aim of being closer to physiological conditions, full-length NCS-1 and a more native-like rRic-8A (residues 1-491) were used. The change in tryptophan fluorescence of rat ΔN31Gα_i1_ upon exchange of GDP for GTPγS was measured both for the intrinsic exchange activity of ΔN31Gα_i1_ and exchange activity following incubation with either rRic-8A-491 or His-NCS-1/rRic-8A-491 in the presence of 0-500 μM CaCl_2_ (Fig S6). Nucleotide exchange rates were determined by fitting data to a single exponential curve, or in the case of rRic-8A-491 not complexed with NCS-1, a double exponential curve in which the slower of these two rates (Fig 8) corresponds to Ric-8A-491-catalyzed nucleotide exchange. The fast phase corresponds to GTPγS binding to an intermediary complex of Ric-8A with nucleotide-free ΔN31Gαi1 that is generated after GDP is released from ΔN31Gα_i1_ upon binding to Ric-8A during incubation of the two proteins. Of note, this intermediary complex was not observed for the NCS-1/Ric-8A complex.

In the absence of Ca^2+^, Ric-8A-491 increases the rate of nucleotide exchange at ΔN31Gα_i1_ almost 4-fold over the intrinsic rate, whereas His-NCS-1/Ric-8A-491 does not significantly affect the exchange rate (Fig 8A). However, the addition of increasing concentrations of CaCl_2_ produces an enhancement of the exchange rate in the presence of His-NCS-1/Ric-8A-491. At 1 μM CaCl_2_, 80 % of the nucleotide exchange rate is restored to levels similar to that observed in the presence of Ric-8A-491 alone (Fig 8). Thus, NCS-1 complexed to Ric-8A effectively inhibits the GEF activity of Ric-8A whereas addition of CaCl_2_ appears to restore GEF activity. From these data, it is possible to calculate a Ca^2+^ activation apparent constant (K_a_ = 0.19 ± 0.23 μM; Fig 8B) for its action at the NCS-1/Ric-8A complex, which disassembles upon Ca^2+^ binding to NCS-1 EF-4, enabling Ric-8A to catalyze Gα_i1_ nucleotide exchange. At this juncture, the effect of CaCl_2_ on intrinsic rate of ΔN31Gα_i1_ nucleotide exchange is not understood.

## Discussion

Several studies have shown the structural determinants of Ric-8A chaperone and GEF activity for Gα protein subunits. Ric-8A phosphorylation promotes Gα recognition since it stabilizes a conformation that allows Ric-8A to trap the Ras-like domain of Gα (Fig 1B). This facilitates the folding of the Gα subunit and stabilizes a nucleotide-free state, which in turn would prepare the protein for GTP loading (*38–40*). However, the mechanism by which the Gα chaperone and GEF activity of Ric-8A could be downregulated by NCS-1 remained unclear. Here, we have combined biochemical, biophysical and protein-protein interaction assays with X-ray crystallography and cryo-EM to shed light into the mechanism that keeps Ric-8A inactive.

The combination of the crystallographic work using Ric-8A peptides of different lengths, together with cell-based protein-protein interaction assays, has enabled the discovery of the necessary and sufficient region of Ric-8A that is required for NCS-1 recognition, a region that is conserved between human and rat (Fig 2 and 3). This region corresponds to the two-helix bundle that constitutes the HEAT repeat 9 of the ARM-HEAT repeat domain, and a disordered region that, in the presence of Gα, forms an extended coil to permit phosphorylated S435 and T440 (in human S436 and T441) to attach to positively charged patches of Ric-8A (Fig 1B and 7) (*38*–*41*). Binding of Ric-8A to Gα does not substantially alter the conformation of the ARM-HEAT repeat domain, since the overall structure and orientation of repeats of uncomplexed rRic-8A and rRic-8A bound to Gα are very similar, such that any major differences are confined to the polypeptide loop regions. However, the binding of NCS-1 to Ric-8A induces a reorganization of the C-terminus of the ARM-HEAT repeat domain that impacts two repeats: the so-called ARM repeat 8 and the HEAT repeat 9. As shown by the crystal structures presented here, the HEAT repeat 9 undergoes an important conformational change: helix a9 unfolds while helix b9 refolds, resulting in the named regions R1 and R2 (Fig 3B and Fig 7B). The relative orientation of R1 and R2 change with respect to a9 and b9. If helices b9 and R2 are superposed, helix a9 is rotated ∼180 degrees with respect to R1 (Fig 7B). To illustrate the structural rearrangement, a morph movie has been produced starting at the Gα-bound and ending at the NCS-1-bound conformations (Movie S1). The structural reorganization affects the distribution of hydrophobic and positively charged amino acids. In the Ric-8A/Gα complex, hydrophobic residues in the HEAT repeat 9 are facing the ARM repeat 8 to create contacts between the two repeats (Fig 7A, C and Movie S1). In the NCS-1ΔH10/Ric-8A-P complex, the same hydrophobic residues are exposed to the solvent, affording their interaction with the hydrophobic crevice of NCS-1, while positively charged residues are exposed on the opposite face, generating an amphipathic structure (Fig 3C, S2, 7C and Movie S1). This would create electrostatic repulsions with repeat 8 and would force the detachment of repeat 9 from repeat 8, so that NCS-1 properly recognizes Ric-8A. The unfolding of helix a9 could act as a hinge to assist in the rearrangement that occurs when Ric-8A binds NCS-1. The cryo-EM structure of NCS-1ΔH10 bound to rRic-8A-452 at low resolution supports this mechanism and suggests that the structural reorganization additionally affects the ARM repeat 8, specifically the long A8-B8 loop and helices B8 and C8 (residues 356 to 400) (Fig 7C and D). Compared with the Ric-8A/Gα complex, the relative position of NCS-1 with respect to Gα is completely different, due to the structural reorganization of these repeats, showing a spring-like behavior (Fig 7D). This conformational rearrangement and the generation of several hinge regions could account for the heterogeneity of the protein complex and the low resolution of the cryo-EM maps.

**Figure 7:**
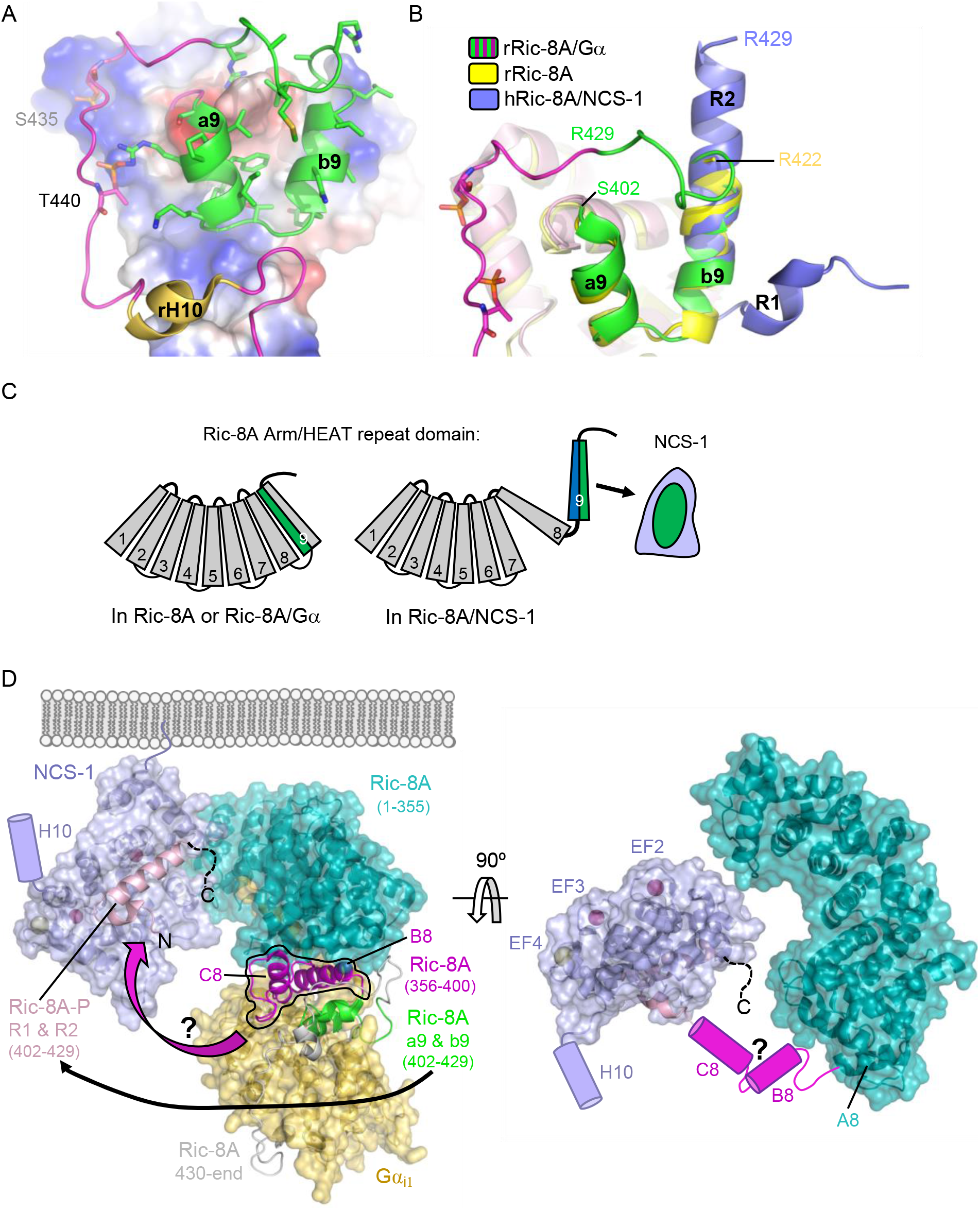
Structural reorganization of Ric-8A for NCS-1 recognition. **A)** The structure of the rRic-8A/Gα_i1_ complex (PDB: 6UKT, (*39*)). Electrostatic potential surface representation of ARM-HEAT domain (repeats 1 to 8). The repeat 9 is shown as ribbons. The Ric-8A region present in the NCS-1/Ric-8A-P crystal structure is shown in green and side chains of the corresponding residues in stick mode. Phosphorylated S435 and T440 are indicated. **B)** Superposition of the structures of Ric-8A bound to Gα (magenta and green, PDB: 6UKT, (*39*)), uncomplexed rRic-8A (yellow, PDB: 6NMG, (*41*)), Ric-8A peptide (light purple) bound to hNCS-1. Ric-8A helix R2 of the complex with NCS-1 was superposed with helix b9 of uncomplexed Ric-8A. **C)** Schematic representation of Ric-8A ARM/HEAT repeat domain (repeats 1 to 9 are indicated) explaining the detachment of repeats 8 and 9 for NCS-1/Ric-8A assembly. The redistribution of charged (blue) and hydrophobic residues (green) in Ric-8A repeat 9 generates the platform for NCS-1 recognition. **D)** Structural comparison between the rRic-8A/Gα complex (PDB: 6UKT, (*39*)) with the NCS-1/Ric-8A X-ray/cryo-EM model presented in this work. Ric-8A ARM-HEAT domains (from repeat 1 to repeat 8 helix A8) have been superposed. The helices B8 and C8 from repeat 8 (magenta) and a9 and b9 from repeat 9 (bright green), which rearrange for NCS-1 recognition, are shown in ribbons. Ric-8A helices a9 and b9 (complex with Gα_i1_) convert to helices R1 and R2 (light pink) when interacting with NCS-1. A magenta curved arrow represents the conformational change that Ric-8A repeat 8 (356-400) may undergo in order to connect with Ric-8A helices R1 and R2. The NCS-1 C-terminal helix is represented as a cartoon. On the right, a top view of the NCS-1/Ric-8A complex is displayed to shown how NCS-1 Ca^2+^ binding sites are exposed to the solvent, and how Ric-8A ARM repeat 8 helices B8 and C8 (magenta cartoons) would occupy the unassigned extra density observed in the cryo-EM map to connect with Ric-8A helices R1 and R2. A dashed black line represents the C-terminus of Ric-8A which gets trapped between NCS-1 and the Ric-8A ARM-HEAT repeat domain.

The model for the NCS-1/Ric-8A complex obtained from the high-resolution crystal structure of hNCS-1ΔH10/Ric-8A-P combined with the low resolution cryo-EM structure of NCS-1ΔH10/rRic-8A-452 meets several criteria that account for the function of a calcium sensor that is anchored to the membrane. The orientation that NCS-1 adopts in the complex leaves the Ca^2+^ binding sites exposed to the solvent for ion exchange, and very importantly, the relative position of NCS-1 and Ric-8A enables membrane anchorage of the Ca^2+^ sensor through its myristoylated N-terminus (*49–51*) (Fig 7D). The NCS-1 dynamic helix H10, which is not present in the construct used for our structural studies, could be exposed to the solvent (Fig 7D). Finally, there is no EM density for the Ric-8A region comprising residues 430-452, which must be disordered and oriented towards the concave face of Ric-8A, sandwiched between NCS-1 and the Ric-8A ARM-HEAT repeat domain (Fig 7D). Indeed, these residues are disordered in the crystal structure of Ric-8A-452 (*41*).

The dynamic NCS-1 C-terminal helix H10 regulates the binding of Ric-8A by inserting into the hydrophobic crevice (Fig 1A) (*25*). Comparison of the structure of Ric-8A-P in complex with NCS1ΔH10 with that of NCS-1 in which its helix H10 is inserted in the hydrophobic crevice explains why this helix constitutes a built-in competitive inhibitor of the NCS-1/Ric-8A interaction (Fig S7A). The helix H10 completely overlaps with the Ric-8A region that is recognized by NCS-1 and displaces most of the H-bonds and contacts reported for the complex, including the C-terminus of helix R1, the N-terminus of helix R2 and the loop connecting them (Fig S2B, C and S6A). Interestingly, the binding of Ric-8A does not cause significant structural reorganization of NCS-1. Superimposition of the hNCS1ΔH10/Ric-8A-P structure with that of free human NCS-1 (*27*) shows subtle changes: the NCS-1 helix H3 and the loop connecting the helices H3 and H4 rearrange to open up the cavity to accommodate helix R2 (Fig 8C). In addition, the loop connecting helices H7 and H8 helices, which interacts with the Ric-8A R1 region, undergoes a conformational change to permit contacts between NCS-1 T135 with Ric-8A residues I407 and, to a lesser extent, V403 (Fig S2A, S2D and Fig S7C).

**Figure 8:**
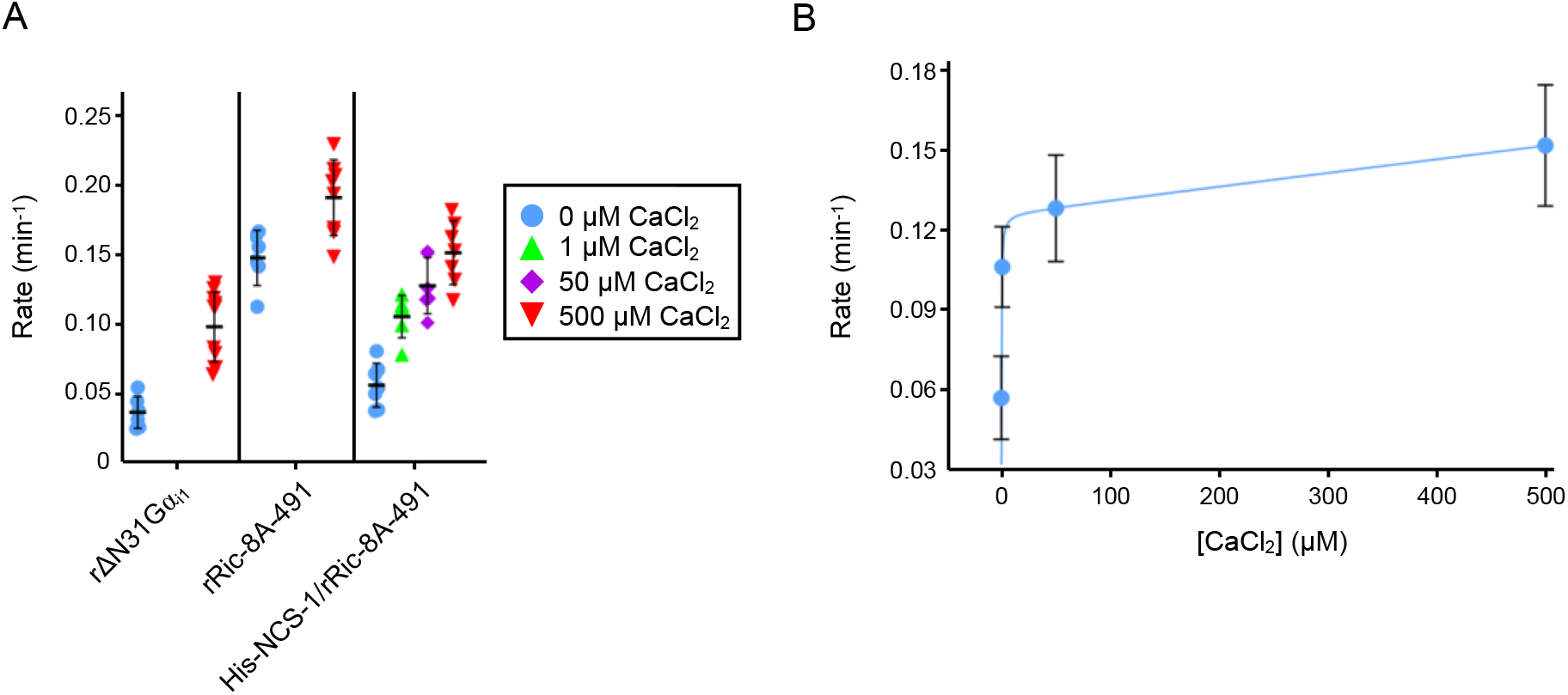
Effect of full-length NCS-1 on GEF activity of rRic-8A-491. **A)** GTP binding rates were measured by following the increase in rat ΔN31Gα_i1_ tryptophan fluorescence following addition of 10 μM GTPγS. Prior to GTPγS addition, ΔN31Gα_i1_ (1 μM final concentration) was incubated with either rRic-8A-491 or His-NCS-1/rRic-8A-491 (0.5 μM final concentration) and 0, 1, 50 or 500 μM CaCl_2_ for five minutes in 50 mM HEPES pH 8, 200 mM NaCl, 2 mM MgCl_2_, 1 mM TCEP before addition of GTPγS. Dunnett’s multiple comparisons test probabilities that nucleotide exchange rates fall within the distribution of ΔN31Gα_i1_ (0 μM CaCl_2_, *n*=6) are: ΔN31Gα_i1_ (500 μM CaCl_2_, *n* = 11), *p* = 0.0096; rRic-8A-491 (0 μM CaCl_2_, *n*=6), *p* = 0.0004; rRic-8A-491 (500 μM CaCl_2_, *n*=8), *p* = 0.0002; His-NCS-1/rRic-8A-491 (0 μM CaCl_2_, *n*=7), *p* = 0.2925 (not significant); His-NCS-1/rRic-8A-491 (1 μM CaCl_2_, *n*=6), *p* = 0.0009; His-NCS-1/rRic-8A-491 (50 μM CaCl_2_, *n*=6), *p* = 0.0007; His-NCS-1/rRic-8A-491 (500 μM CaCl_2_, *n*=7), *p* = 0.0003. In all cases means and standard deviations are reported for a minimum of 6 experimental replicates. **B)** GTP binding rates of His-NCS-1/rRic-8A-491 plotted vs CaCl_2_ concentration. Data were fit to a One-site binding model (*77*).

The data presented here show how Ca^2+^ and Ric-8A phosphorylation serve as determinants of NCS-1/Ric-8A recognition. Binding of Ca^2+^ to the structural EF-hands EF-2 and EF-3 is necessary for protein recognition, while EF-4, the regulatory Ca^2+^ binding site (*15*), is free of Ca^2+^, as shown in the crystal structure of the NCS-1ΔH10/Ric-8A-P complex (Fig 4A) and the *in vitro* reconstruction of the complex (Fig 2A). We show that a fully Ca^2+^ loaded protein does not properly recognize Ric-8A, suggesting that in the cellular context, an increase in Ca^2+^ concentration and subsequent loading of EF-4 may trigger the disruption of the NCS-1/Ric-8A complex. This agrees with previous cell-based protein-protein interaction assays that showed weakened affinity of NCS-1 for Ric-8A in Ca^2+^ saturating conditions (*25*). Nucleotide exchange assays demonstrate a loss of Ric-8A GEF function when in complex with NCS-1, while nucleotide exchange rates are rescued following addition of Ca^2+^ (Fig 8). With respect to the phosphorylation state of Ric-8A, NCS-1 binds to unphosphorylated Ric-8A protein (Fig 2A and 6A) and more importantly, in the context of the protein complex, NCS-1 protects Ric-8A from CK2-mediated phosphorylation (Fig 6B). In fact, the structure of the NCS-1/Ric-8A complex obtained with the combination of the crystallographic and cryo-EM data explains the structural determinants of NCS-1 protection, showing that the phosphorylatable residues S335 and T440 would remain completely occluded between NCS-1 and the concave face of the Ric-8A ARM-HEAT repeat domain, and therefore, unavailable to CK2 (Fig 7D). In this context, an increase of Ca^2+^ levels would permit the disassembly of the NCS-1/Ric-8A complex, allowing release of Ric-8A from the membrane, and subsequent phosphorylation (Fig 9). This would promote Ric-8A activation for Gα subunit binding and GEF activity. When inactivation of Ric-8A is required in the cellular context, Ric-8A dephosphorylation must precede NCS-1 binding, since the phosphorylated protein does not recognize NCS-1 properly (Fig 6C and 9). This can be explained as well in the context of the structure of phosphorylated Ric-8A bound to Gα, since the strong ionic interactions of phosphorylated S335 and T440 with the ARM-HEAT repeat domain would preclude the structural rearrangement of repeats 8 and 9 (Fig 7A). Cellular mechanisms of Ric-8A dephosphorylation are not understood. It has been proposed that in the cell, Ric-8A is constitutively phosphorylated (*32*). However, and in the light of the results presented here, in neurons and other tissues where NCS-1 is expressed (*2*), Ric-8A phosphorylation may be under NCS-1 control and regulated by Ca^2+^ signals. Since NCS-1 is a Ca^2+^ sensor that is constantly bound to the membrane and close to Ca^2+^ channels, changes of cytosolic Ca^2+^ would promote the disassembly of the protein complex and the subsequent phosphorylation of Ric-8A by CK2 (Fig 9). Given that Ric-8A is a ubiquitously expressed protein, it is unknown if, in tissues where NCS-1 is not expressed, there are other Ca^2+^ sensors in charge of the regulation of its activity. Furthermore, it is likely that NCS-1 interacts with Ric-8B, a chaperone for other Gα (*29, 30*), since the sequence of the NCS-1 interacting region is conserved.

**Figure 9:**
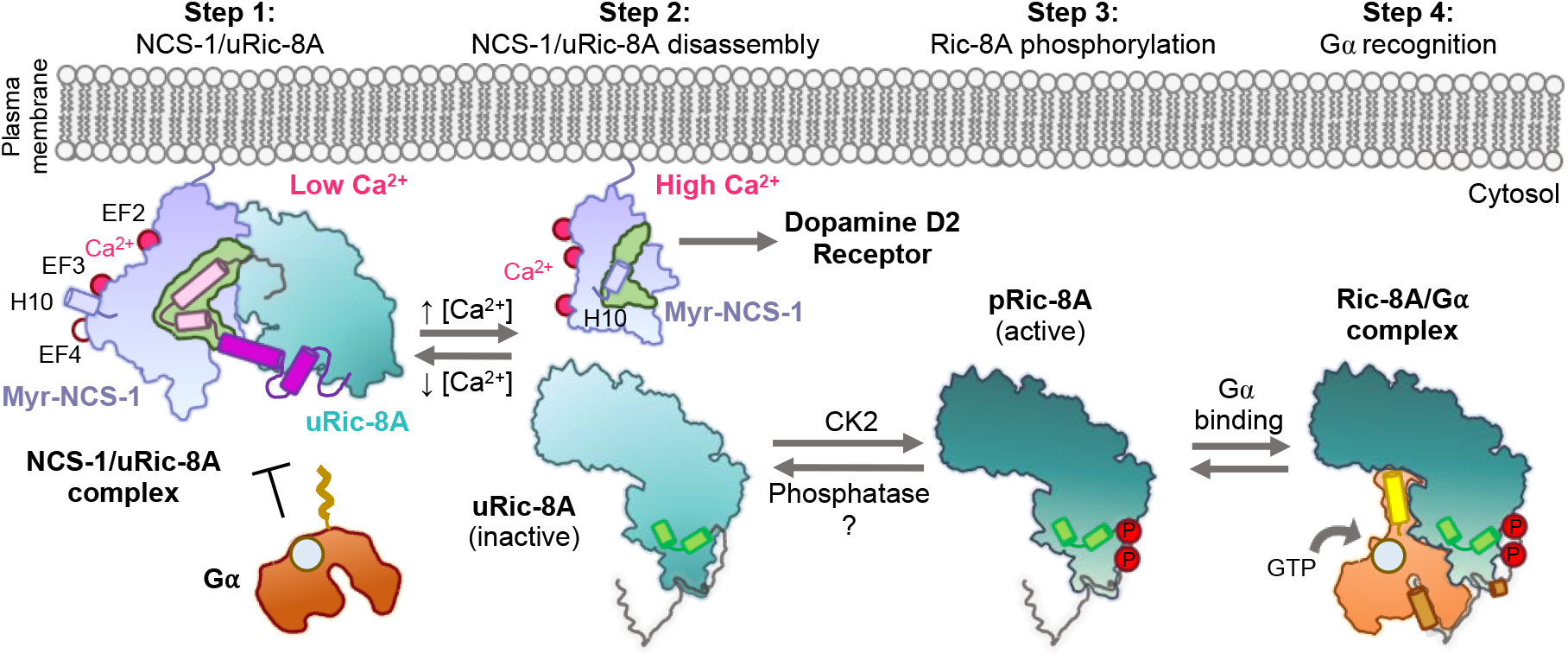
Schematic representation of the mechanism of Ric-8A activation regulated by NCS-1. Step 1: At low Ca^2+^ concentrations NCS-1 interacts with unphosphorylated Ric-8A (uRic-8A), at the plasma membrane. NCS-1 protects Ric-8A from phosphorylation or Gα subunit binding by inducing a conformational rearrangement of repeats 8 (helices B8 and C8 in magenta) and 9 (helices R1 and R2 in pink). Step 2: When Ca^2+^ levels increase in the cytosol, NCS-1 binds Ca^2+^ at EF-4 and the complex is disassembled. NCS-1 helix H10 inserts in the hydrophobic crevice (green) and would be ready for Dopamine D2 receptor recognition. Inactive uRic-8A, free of NCS-1, repacks repeats 8 and 9 (helices a9 and b9 in light green) and S335 and T440 are phosphorylated (P) by CK2 (Step 3). In this state, phosphorylated Ric-8A (pRic-8A) is now active, recognizes prefolded Gα subunit and allows GTP loading (Step 4).

The transmission of the nerve impulse between neurons begins with the generation of an action potential in the axon initial segment. An action potential is a depolarization of the plasma membrane that depends on Na^+^ entry through specific voltage-gated channels and that propagates along the axon to the terminal. When the depolarization Na^+^ wave reaches the axon terminal, voltage dependent Ca^2+^ channels open allowing the massive entrance of Ca^2+^ which in turn triggers neurotransmitter release (*52, 53*). Therefore, Na^+^ precedes Ca^2+^ waves. Here we demonstrate that Na^+^ decreases the affinity of NCS-1 for Ca^2+^ (Fig 5D and Table 2). The crystal structure of the hNCS-1ΔH10/Ric-8A-P complex shows that NCS-1 binds Na^+^ at EF-4, the regulatory Ca^2+^ binding site. From a physiological point of view this finding is relevant since NCS-1/Ric-8A disassembly and Ric-8A activation would occur at a Ca^2+^ signal that is higher than initially expected, ensuring the return to the Ric-8A inactive and NCS-1-complexed state when Ca^2+^ levels decrease, due to sequestering by several high capacity, low specificity, Ca^2+^ binding proteins.

This work also sheds light into the mechanism of specificity that allows NCS-1 to recognize different targets. NCS-1 interacts with several proteins related to G-protein signaling: Ric-8A and GPCRs such as the adenosine A2A, Cannabinoid CB1 or the Dopamine D2 receptor (*21, 22, 24*). The structure and Ca^2+^ determinants of Dopamine D2 receptor recognition by NCS-1 are known (Fig 1) (*23*). It is relevant that the Ca^2+^ signals that trigger protein-protein recognition are opposite: while the binding of NCS-1 to Ric-8A takes place at low Ca^2+^ concentrations at which the functional site EF-4 is free of Ca^2+^ (Fig 3A and 4A), binding to Dopamine D2 receptor takes place at high Ca^2+^ levels (Fig 1A). Therefore, while Ric-8A is being blocked by NCS-1, Dopamine D2 receptor is free of NCS-1 and *vice versa*. Also, the role of the NCS-1 dynamic C-terminal helix H10 is different in the two molecular recognition processes, while NCS-1 uses this helical element to properly recognize Dopamine D2 receptor (*23*), the helix H10 negatively regulates the binding to Ric-8A. Finally, if the crystal structure of the NCS-1ΔH10/Ric-8A-P complex is compared with other NCS proteins bound to their corresponding targets, the complex that is structurally most similar is the KChIP1/Kv4.3 pair, with an elongated helix resembling the R2 bound to Ric-8A (Fig S7D). Interestingly, the N-to C- orientation of the helix is opposite, and no extra helical segment (R1) is recognized. In addition, the Ca ^2+^ content is different (Fig 1A) (*54*). The number of structures that have been determined of NCS proteins bound to different targets continues to grow. Comparison of these structures show that, while all of the ligands use the same NCS hydrophobic crevice, differences in Ca^2+^ site ligation, role of the helix H10 and the loop that connects EF-3 and EF-4 in NCS-1 shapes the crevice to recognize, very specifically, different target protein motifs. In this sense polar interactions between NCS-1 and the target protein play also an important role to ensure specificity and affinity (Fig 1A and S7).

Small-molecule PPI modulators with therapeutic potential have been discovered in the past few years (*6, 27, 55*). They are able to inhibit (e.g. the phenothiazine FD-44) or stabilize (e.g. the acylhydrazone 3b) the formation of the NCS-1/Ric-8A complex and in doing so, they regulate synapse number and function and show a promising prospect in the treatment of neurodevelopmental disorders (*6, 26*) and neurodegenerative diseases (*27*). It has been suggested that these small molecules, which bind to the same NCS-1 site despite their opposite activity, are not Ric-8A competitors. Instead, they have been suggested to be allosteric modulators that contribute to the stabilization of the dynamic helix H10 inside (inhibitors) or outside (stabilizers) the crevice, favoring NCS-1 conformations that hinder or allow the entrance of Ric-8A and the subsequent formation of the protein complex. Supporting this, the modulator FD-44 does not inhibit the formation of the NCS-1/Ric-8A complex when the dynamic helix H10 is deleted. The structure of the hNCS-1ΔH10/Ric-8A-P complex explains the molecular mechanism of action of the regulatory compounds. Comparison of the different complexes shows that the binding sites for these molecules overlap with the C-terminal half of helix R2 (Fig S7A and S7B). With respect to the inhibitors, hydrophobic interactions of FD-44 with the helix H10 stabilize the crevice-inserted helix H10 conformation of NCS-1. The combination of the compound plus helix H10 makes the Ric-8A interacting region of NCS-1 completely unavailable (Fig S7A). On the contrary, for the PPI stabilizer, the polar characteristics of the molecule hinders the approach of the helix H10 to the crevice, making the C-terminal part of the cavity, which is more relevant in terms of the PPI, available for Ric-8A recognition (Fig S7B). The structure of hNCS-1ΔH10/Ric-8A-P does not support the existence of a ternary NCS-1/Ric-8A/stabilizer complex, since there would be no room for the repositioning of the compound. Therefore the binding of Ric-8A to NCS-1 would displace modulator 3b from the crevice, which would be likely given the moderate affinity of the compound (*27*). Finally, we believe that the high-resolution data on the NCS-1/Ric-8A PPI interface presented here will be essential to rationally develop improved compounds with better PPI regulatory properties and selectivity, since the relevant interactions between NCS-1 and Ric-8A have been finally revealed. The fact that the NCS-1 Ca^2+^ and structural determinants are different in recognizing Ric-8A and the Dopamine D2 receptor is conceptually relevant, since opens the path to the design of selective drugs that target specifically these neuronal pathways.

## Material and methods

### Cloning, expression and purification of proteins

*NCS-1 constructs:* Human full-length NCS-1 (100 % protein sequence identity with the rat variant) was cloned in the pETDuet vector (*27*). A stop codon was introduced after residue P177 to generate the NCS-1ΔH10 construct and using the IVA cloning strategy (*56, 57*). The proteins were overexpressed in *E. coli* (BL-21*) as reported (*27, 58*). Briefly, cells were resuspended in lysis buffer (50 mM Hepes pH 7.4, 100 mM KCl, 1 mM 1,4-dithiothreitol (DTT), 0.1 mM phenylmethylsulfonyl fluoride (PMSF), 10 μg/ml DNAse) and disrupted by sonication. The lysate was centrifuged at 16,000 rpm in a SS-34 rotor (4 ºC, 45 min). 1 mM CaCl_2_ was added to the clarified supernatant and the resulting solution was injected into a Hi Trap Phenyl FF hydrophobic column (Cytiva) preequilibrated with lysis buffer plus 1 mM CaCl_2_. The column was washed with 5 column volumes of HIC-A buffer (20 mM Tris pH 7.9, 1 mM CaCl_2_, 1 mM DTT). Protein elution was achieved applying a EGTA gradient with HIC-B buffer (20 mM Tris pH 7.9, 2 mM EGTA, 2 mM DTT). NCS-1ΔH10 elution occurred at 1.2 mM EGTA, while full-length NCS-1 elution occurred at 1.4 mM EGTA. Protein quality was evaluated by nano-DSF (see below) and SDS-PAGE. Fully Ca^2+^-loaded protein was prepared by dialyzing the EGTA-eluted sample against HIC-A buffer. Then, the sample was loaded in an anion exchange HP Q column (Cytiva) and eluted with a NaCl gradient using QB buffer (20 mM Tris pH 7.9, 1 mM CaCl_2_, 500 mM NaCl, 1 mM DTT). NCS-1 elution occurred at 175 mM NaCl.

N-terminally hexahistidine-tagged full-length human NCS-1 (His-NCS-1) was expressed in a pET28a+ vector using BL21(DE3) pLysS *E. coli*. Overnight cultures were grown in 100 ml LB media containing 100 μg/ml kanamycin at 200 rpm and 37 °C. After ∼16 h cells were pelleted at 2200 x g for 10 min at 4 °C using a benchtop Sorvall Legend RT. Re-suspended pellets were added to 1 L 2xYT media containing 100 μg/ml kanamycin and incubated at 37 °C and 200 rpm until an OD_600_ of 0.8-1 was achieved and cells were induced with 0.3 mM isopropyl β-D-1-thiogalactopyranoside (IPTG) at 16 °C. Approximately 18 hours post-induction, cells were pelleted at 12,000 x g for 15 min in a Sorvall RC6+ Centrifuge and cell pellets were stored at -80 °C. Pellets were resuspended in 50 ml (per L of cell pellet) of 50 mM HEPES pH 7.4, 100 mM KCl, 2 mM beta-mercaptoethanol, 10 μg/ml DNase and lysis was performed with an Avestin Emulsiflex-C5 homogenizer. Lysate was cleared for 40 min at 4 °C and 35,000 rpm with a Beckman Coulter Optima XE-90 ultracentrifuge and a Type 45 TI rotor and the supernatant was loaded on a HisTrap FF crude column. NCS-1 eluted at 25 % during a 0-100 % gradient to 50 mM Tris pH 8, 300 mM Imidazole, 1 mM CaCl_2_, 2 mM beta-mercaptoethanol. Eluted His-NCS-1 was dialyzed overnight in HIC-A buffer and dialyzed His-NCS-1 was loaded on a 5 ml HiTrap Phenyl HP column and eluted, as described above, in HIC-B buffer.

#### Ric-8A constructs

Rat Ric-8A-452 (residues 1-452) was previously cloned in pET28a vector (*59*). Introduction of stop codons after residues G423 and G432 resulted in the so-called Ric-8A-423 and Ric-8A-432 constructs. Proteins were expressed in Rosetta2 pLysS cells and purified similarly to previously reported methods (*41*). Briefly, cells were resuspended in lysis buffer (50 mM Tris pH 8, 250 mM NaCl, 20 mM imidazole, 5 % glycerol, 2 mM beta-mercaptoethanol) with 0.1 mM PMSF, 10 μg/ml DNAse and protease inhibitors (cOmplete, EDTA-free cocktail, Roche), and disrupted with a sonicator. After centrifugation (45 Ti rotor at 30,000 rpm and 4 ºC for 40 minutes) the clarified supernatant was loaded in a Nickel-affinity column (HisTrap FF, Cytiva). The column was washed with 10 volumes of lysis buffer and protein elution was achieved with a gradient with NiB buffer (50 mM Tris pH 8, 250 mM NaCl, 500 mM imidazole, 5 % glycerol, 2 mM beta-mercaptoethanol), after an extra wash step with 7 % NiB buffer. Protein sample was next dialyzed in GF buffer (50 mM Tris pH 8, 150 mM NaCl, 5 % glycerol, 2 mM beta-mercaptoethanol) for imidazole removal. Tobacco Etch Virus (TEV) protease was used to cleave the hexahistidine-tag off using a 1:40 molar ratio (TEV:His-Ric-8A) during 16 h. After TEV treatment, the sample was incubated with Ni^2+^-chelated Sepharose HP beads (Cytiva) to get rid of the uncleaved protein. A final polishing step was performed in a size exclusion column (HiLoad 16/600 Superdex 200 pg, Cytiva) preequilibrated in GF buffer. Protein quality was evaluated by nano-DSF (see below) and SDS-PAGE.

N-terminally hexahistidine-tagged Rat Ric-8A residues 1-491, Ric-8A-491, was expressed and purified as previously described in a pET28a vector with BL21(DE3) pLysS *E. coli* (*41, 59, 60*). Post hexahistidine-tag removal by TEV protease and purification by Source 15Q column (*39*), Ric-8A-491 was buffer exchanged into 50 mM Tris pH 8, 250 mM NaCl, and 1 mM Tris(2-carboxyethyl) phosphine (TCEP) for assembly of protein complex.

#### Gα construct

N-terminally glutathione-S-transferase tagged rat Gα_i1_ with a 31 residue N-terminal truncation (ΔN31Gα_i1_) was expressed from a pDest15 vector in BL21(DE3) RIPL *Escherichia coli* and purified as previously described (*60, 61*).

#### Ric-8A nanobodies

Specific rat Ric-8A nanobodies Nb8109, Nb8117 and Nb8119 were expressed and purified as reported (*39*) and concentrated to 10 mg/ml in 50 mM HEPES pH 8, 150 mM NaCl and 1 mM TCEP for complex assembly.

### Assembly of protein complexes

#### NCS-1ΔH10/rRic-8A-452 complexes

##### Ca^2+^ free conditions (Assembly (i))

Pure rat His-Ric-8A-452 (50 mM Tris pH 8, 250 mM NaCl, 5 % glycerol, 2 mM beta-mercaptoethanol) was mixed with NCS-1ΔH10 (20 mM Tris pH 7.9, 1.2 mM EGTA, 1 mM DTT) in a 1:1.9 (Ric-8A:NCS-1) molar ratio. Final NaCl and EGTA concentrations were adjusted to 125 mM and 0.6 mM, respectively. To ensure absence of Ca^2+^, the mixture was dialyzed against buffer 50 mM Tris pH 8, 200 mM NaCl, 2 mM EGTA, 1 mM TCEP, with and without 1 mM MgCl_2_ (2 changes, first after 4 h and second for 16 h). The final sample was concentrated and subjected to a size exclusion chromatography in the same buffer and using a Superdex 200 HR 10/30 column (Cytiva). 12 % SDS-PAGE gels were run to identify the composition of eluted samples.

##### High Ca^2+^ conditions (Assembly (ii))

Protein mixture was performed as above but the purified fully Ca^2+^ loaded NCS-1ΔH10 (20 mM Tris pH 7.9, 2 mM CaCl_2_, 1 mM DTT) was used instead. The final NaCl and CaCl_2_ concentrations in the protein mixture were 125 mM and 2 mM, respectively. After 1h incubation, the sample was concentrated and subjected to a size exclusion chromatography. as described above, but the column was equilibrated in buffer 50 mM Tris pH 8, 200 mM NaCl, 2 mM CaCl_2_, 1 mM TCEP.

##### Dialysis from EGTA to Ca^2+^ conditions (Assembly (iii))

Protein mixture was the same as that for Ca^2+^ free conditions and the resulting sample (at 0.6 mM EGTA) was dialyzed against buffer containing 50 mM Tris pH 8, 200 mM NaCl, 2 mM CaCl_2_, 1 mM TCEP. This assembly was performed with both the un-phosphorylated and phosphorylated variants of His-Ric-8A-452. The same assembly was also performed in a K^+^ containing buffer. For this, unphosphorylated His-Ric-8A-452 was first dialyzed against buffer containing 50 mM Tris pH 8, 250 mM NaCl, 5 % glycerol, 2 mM beta-mercaptoethanol, to replace NaCl for KCl. Protein mixture, dialysis and gel filtration were carried out substituting NaCl for KCl.

#### NCS-1ΔH10/rRic-8A-452/Nb8109/Nb8117/Nb8119 complex

Purified Ric-8A nanobodies (Nbs) 8109, 8117 and 8119 were added to the freshly reconstituted NCS-1ΔH10/rRic-8A-452 complex in a molar ratio Nb:NCS-1/Ric-8A 1.6:1 and incubated for 16 h. Size exclusion chromatography was run with a Superdex 200 HR 10/30 column to remove the excess of non-assembled nanobodies. The presence of the five components in the final complex was assessed by MALDI-TOF spectrometry. Furthermore, SEC-MALS assays confirmed the purification of a homogeneous complex with an estimated molecular mass of 117-121 kDa, in agreement with the calculated mass of 115 kDa.

### Assembly, crystallization, diffraction data collection and structure solution of hNCS-1ΔH10/Ric-8A-P complexes

Three highly pure (>95 %) HPLC-verified Ric-8A peptides were purchased from GenicBio for structural studies. They ranged from residue 400 to residues 423 (P1), 429 (P2) and 432 (P3) (Fig 1B). Lyophilized peptides were solubilized in HIC-B buffer and mixed with purified NCS-1ΔH10 in a 1:10 (protein:peptide) molar ratio (final EGTA concentration 1.7 mM). The mixture was dialyzed against a buffer containing 20 mM Tris pH 8, 0.5 mM CaCl_2_, 0.5 mM DTT (2 changes, first after 4 h, second for 16 h). Thermal stability of the final samples was evaluated by nano-DSF. The final sample was concentrated to 20 mg/ml with a Vivaspin 2 device (2 kDa cutoff, Sartorius).

Crystallization screenings were set with an Oryx8 robot (Douglas Instruments) at 4 ºC, using the sitting drop vapor diffusion method and mixing equal volumes of protein complex and precipitant. Initial crystals were obtained with P2 and P3 peptides and solution from JBScreen Classic (Jena Bioscience) and INDEX (Hampton Research) crystallization screenings. Diffracting crystals obtained with P2 peptide grew using microseeding techniques and precipitant solution 25 % PEG 4000, 100 mM NaAc pH 5 and 100 mM MgCl_2_. Crystals with peptide P3 grew in two different conditions: 30 % PEG 4000, 100 mM NaAc pH 4.6, 100 mM MgCl_2_ and 25 % PEG 3350, 100 mM NaAc pH 4.5. Crystals were cryo-protected adding 30 % (v/v) glycerol to the precipitant solutions and flash-frozen in N_2_(l).

Diffraction data were collected at 100 K and 0.979 Å wavelength at ALBA synchrotron radiation source (BL13 beamline) (Table 1). Data were processed with AutoPROC using the extended anisotropic method (*62*). The first structure was solved by molecular replacement with Phaser (*63*), using with data from P2 peptide crystals (Structure 1). As search model, the structure of hNCS-1 (PDB: 6QI4), lacking the C-terminal helix H10, was used (*27*). Successive cycles of automatic refinement with Phenix (*64*) and manual building with Coot (*65*) were performed. The refined structure was used to solve Structure 2 and 3, using Fourier differences calculations. The final models were validated with Molprobity (*66*). Details on data processing and refinement are shown in Table 1. The structures were analyzed using different programs from the CCP4 package (*67*) and the PISA server (*43*). Images were prepared with PyMOL (*68*). The final structures were deposited in the PDB with codes: Structure 1 (8ALH), Structure 2 (8AHY), Structure 3 (8ALM).

### Thermal shift assay

Label-free thermal shift assays with hNCS-1 full-length, hNCS-1ΔH10, rRic-8A-452, NCS-1ΔH10/rRic-8A-452, NCS-1ΔH10/Ric-8A-P2 peptide and NCS-1ΔH10/Ric-8A-P3 peptide were performed using a Tycho NT.6 instrument (NanoTemper Technologies). Proteins at 10 μM in their corresponding final buffers (see above) were measured in duplicates using NanoTemper capillaries. Intrinsic fluorescence was recorded at 330 nm and 350 nm while heating the sample from 35 to 95 ° C at a rate of 30 ºC/min. The fluorescence ratio (350/330 nm) and the inflexion temperature, Ti, were calculated by Tycho NT.6 software.

### Phosphorylation assays

Purified rRic-8A-452 and NCS-1ΔH10/rRic-8A-452 were phosphorylated with casein kinase II (CK2, New England Biolabs) as previously described by McClelland et al. (2020) (*39*). Briefly, 2 mg of each were dialyzed (2 changes, 2 h and o/n, 4 ºC) in prephosphorylation buffer (50 mM Tris pH 8, 150 mM NaCl, 2 mM CaCl_2_, 1 mM TCEP). Samples were mixed 1:1 in 2x reaction buffer (100 mM Tris pH 8, 200 mM NaCl, 20 mM MgCl_2_, 2 mM EGTA, 1 mM DTT). Half of the samples was subjected to phosphorylation by adding 300 U CK2 and 5 mM ATP. Reactions were allowed to proceed for 16 h and 18 ºC. The other half of samples were treated similarly but CK2 and ATP were not added (non-phosphorylated sample; controls).

To distinguish between phosphorylated and non-phosphorylated proteins, anion exchange chromatography was performed with the non-phosphorylated samples (controls) and those subjected to CK2 treatment. Samples were dialyzed in RV buffer (50 mM Tris pH 8, 125 mM NaCl, 1 mM CaCl_2_, 1 mM DTT) and injected into an anion exchange HiTrap Q HP column (Cytiva) preequilibrated with QA buffer (50 mM Tris-HCl pH 8, 75 mM NaCl, 1 mM CaCl_2_, 1 mM DTT). Protein elution was achieved with a gradient using QB buffer (50 mM Tris-HCl pH 8, 500 mM NaCl, 1 mM CaCl_2_, 1 mM DTT). Fractions from each peak were collected and analyzed by SDS-PAGE. Presence of phosphorylation in the NCS-1ΔH10/rRic-8A-452 complex was additionally verified in a phosphoprotein assay by LC-MS/MS.

### Phosphoprotein analysis by LC-MS/MS

The phosphoprotein assay was divided into three different steps: a) in-gel sample digestion; b) phosphopeptide purification; and c) protein identification by tandem mass spectrometry.

#### In-gel sample digestion

Gel band samples from 1D gel separation and Coomassie staining, were automatically in-gel digested. Gel bands were excised, cut into cubes (1 mm^2^), deposited in 96-well plates and automatically processed in an OT-2 digestor (Opentrons, New York, USA). The digestion protocol used was based on Shevchenko et al. 1996 (*69*) with minor variations: gel plugs were washed first with 50 mM ammonium bicarbonate and secondly with acetonitrile, prior to reduction and alkylation (5 mM tris(2-carboxyethyl)phosphine) and 10 mM chloroacetamide in 50 mM ammonium bicarbonate solution, at 56 ºC for 30 min). Gel pieces were then rinsed first with 50 mM ammonium bicarbonate, and secondly with acetonitrile, and then were dried under a stream of nitrogen. Pierce MS-grade trypsin (Thermo-Fisher Scientific, Massachusetts, USA) was added at a final concentration of 16 ng/μl in 50 mM ammonium bicarbonate solution, and the digestion took place at 37 °C for 2 h. Peptides were recovered in 50 % ACN/ 0.5 % FA, dried in speed-Vac and kept at -20 °C until phosphopeptide enrichment.

#### Phosphopeptide purification

Phosphopeptide enrichment procedure utilized two concatenated in-house packed microcolumns, immobilized metal affinity chromatography and Oligo R3 polymeric reversed-phase that provided selective purification and sample desalting prior to LC-MS/MS analysis, and was performed as previously reported (*69*).

#### Protein identification by tandem mass spectrometry (LC–MS/MS Exploris 240)

The peptide samples were analyzed on a nano liquid chromatography system (Ultimate 3000 nano HPLC system, Thermo Fisher Scientific) coupled to an Orbitrap Exploris 240 mass spectrometer (Thermo Fisher Scientific). Samples (5 μl) were injected on a C18 PepMap trap column (5 μm, 100 μm I.D. x 2 cm, Thermo Scientific) at 20 μl/min, in 0.1 % formic acid in water, and the trap column was switched on-line to a C18 PepMap Easy-spray analytical column (2 μm, 100 Å, 75 μm I.D. x 50 cm, Thermo Scientific). Equilibration was done in mobile phase A (0.1 % formic acid in water), and peptide elution was achieved in a 30 min gradient from 4 – 50 % B (0.1 % formic acid in 80 % acetonitrile) at 250 nl/min. Data acquisition was performed using a data-dependent top-15 method, in full scan positive mode (range of 350 to 1200 m/z). Survey scans were acquired at a resolution of 60,000 at m/z 200, with Normalized Automatic Gain Control (AGC) target of 300 % and a maximum injection time (IT) of 45 ms. The top 15 most intense ions from each MS1 scan were selected and fragmented by Higher-energy collisional dissociation (HCD) of 28. Resolution for HCD spectra was set to 15,000 at m/z 200, with AGC target of 75 % and maximum ion injection time of 80 ms. Precursor ions with single, unassigned, or six and higher charge states from fragmentation selection were excluded.

MS and MS/MS raw data were translated to mascot general file (mgf) format using Proteome Discoverer (PD) version 2.5 (Thermo Fisher Scientific), and searched using an in-house Mascot Server v. 2.7 (Matrix Science, London, U.K.) against an in-house database including Ric-8A protein sequence along with common laboratory protein contaminants. Search parameters considered fixed carbamidomethyl modification of cysteine, and the following variable modifications: methionine oxidation, phosphorylation of serine/threonine/tyrosine and deamidation of asparagine/glutamine. Peptide mass tolerance was set to 10 ppm and 0.02 Da, in MS and MS/MS mode, respectively, and 3 missed cleavages were allowed. The Mascot confidence interval for protein identification was set to ≥ 95 % (P < 0.05) and only peptides with a significant individual ion score of at least 30 were considered.

### Binding of NCS-1 to Na^+^, K^+^ and Ca^2+^

#### Intrinsic florescence titration assay

Tryptophan emission fluorescence of EGTA-purified full-length NCS-1 was recorded at 10 μM in buffer containing 20 mM Tris pH 8, 100 μM EGTA, 1 mM DTT and 0-300 mM NaCl or 0-300 mM KCl. Data were acquired with a Tycho NT.6 equipment (NanoTemper Technologies). The emission fluorescence intensity was recorded at 330 nm and 35 ºC. Fluorescence intensities were normalized as (I_0_ − I)/I_0_. Three independent experiments were performed. Mean and SEM values were represented at different Na^+^ and K^+^ concentrations. The apparent dissociation constant was calculated by using a least squares algorithm to fit the experimental data to a 1:1 stoichiometry model (*6*). The fitting was performed with KaleidaGraph Data Analysis Program (*70*).

#### Isothermal titration calorimetry (ITC)

Ca^2+^ binding was characterized by ITC at 25 °C using a VP-ITC microcalorimeter (GE Healthcare, Northampton, MA) with a cell volume of 1.4619 ml in Na^+^ or K^+^ containing buffers (20 mM Tris pH 7.9, 2 mM EGTA, 150 mM NaCl or 150 mM KCl). Before measurements, EGTA-purified full-length hNCS-1 (25 μM) was dialyzed in parallel against the above buffers (3×300 ml; 2h, 2h and 20 h) and then against the same buffers without EGTA. Protein solutions at 110 μM were loaded into the calorimetric cell and titrated by stepwise injections of a 1.5 mM CaCl_2_ solution prepared in the final dialysate. Dilution heats, evaluated separately, were found to be negligible. The binding isotherms were fit by nonlinear regression analysis using the AFFINImeter software (*71*) using the model builder to create a sequential binding model with three different binding sites. Binding constants (K_d_) and binding enthalpy changes (ΔH_d_) were directly determined from data fitting. The free energy change was calculated as Δ*G* = −*R*T ln K_d_ (*R* = 1.986 cal mol^-1^ K^-1^) and the entropy change using the Gibbs equation (ΔG = ΔH − T ΔS).

### Co-immunoprecipitations

Human NCS-1 and V5-tagged Ric-8A construct were previously described (*6*). Using the IVA cloning strategy (*56, 57*) deletion constructs were prepared ending at residues G424 (hRic-8A-424, which corresponds to G423 in the rat variant) and G433 (hRic-8A-433, in rat, G432). Furthermore, a full-length hRic-8A mutant (S436A and T441A) was prepared to avoid phosphorylation of the protein at these sites. Constructs were cotransfected into HEK293 cells using Lipofectamin 2000 (Thermo) following manufacturer instructions. 48 h after transfection cells were lysed in lysis buffer (150 mM NaCl, 1.0 % Nonidet P-40, 50 mM Tris pH 8.0). Lysates were then incubated overnight (12 h) at 4 °C with mouse anti–NCS-1 (1:500; Cell Signaling). Samples were subsequently incubated for 2 h with Protein-G–Sepharose (Sigma-Aldrich). After 3 washes with lysis buffer, proteins were eluted from the Sepharose and analyzed by Western blot following standard procedures; 10 % of the lysate before immunoprecipitation was run as input. Mouse anti-V5 (1:2000; Thermo) and rabbit anti–NCS-1 (1:1000; Cell Signaling) antibodies were used for Western blot. The Inmunoprecipitation blot was incubated with anti-mouse TrueBlot (Rockland) as secondary antibody to avoid heavy-/light-chain antibody interference.

### Cryo-Electron Microscopy

For cryo-EM grid preparation 300 mesh Quantifoil 0.6/1 Au grids were glow discharged with a Leica EM Ace200 Vacuum Coater at 15 mA for 60 s prior to adding 3 μl of pure NCS-1 ΔH10/rRic-8A-452/Nb8109/Nb8117/Nb8119 complex at 0.6 mg/ml.

Grids were vitrified with ethane using a Vitrobot Mark IV (FEI Company) and stored in liquid nitrogen until data collection. 9,776 movies were collected in a Titan Krios at 300 kV using a K3 detector at the European Synchroton Radiation Facility (ESRF) (*48*). Movies were recorded at a magnification yielding 0.84 Å/pixel with a dose rate of 16.3 e^-^/Å^2^/s and a defocus range between -1 to -3 μm using the Smart EPU Software (Thermo Fisher Scientific). Movies were built of 50 frames each with a total dose of 50.8 e^−^/Å^2^ (∼1 e^−^/Å^2^ per frame) using a total exposure time of 2.2 s. Data processing was performed using RELION-4.0 (*72*) unless otherwise specified. Drift and beam-induced motion correction (5 × 5 patches) were performed using MotionCor2 (*73*) together with dose weighting. Contrast transfer function (CTF) estimation and determination of defocus range were performed using CTFFIND-4.1 (*74*). Particle picking was carried out with Topaz (*75*), where 3,418,186 particles were extracted and input to two rounds of 2D classifications to remove false-positives. An *ab initio* model was generated followed by five rounds of 3D classification on 886,718 particles. The final model contained 37,543 particles which resulted in a 9.3 Å map following the gold-standard FSC cut-off of 0.143. Since the map did not display secondary structure features, likely due to conformational heterogeneity, we low-pass filtered the map to 17 Å, a resolution which we felt confident that the map was devoid of noise. Docking into the cryo-EM map was performed using the coordinates of rRic-8A (1-400) bound to nanobodies Nb8109, Nb8117 and Nb8119 (rRic-8A-400/Nb8109/Nb8117/Nb8119) from the rRic-8A/Gα_i_/nanobody complex (PDB: 6UKT, docking residues 1-355 for best fit) and the crystal structure of hNCS-1/Ric-8A-P determined in the current work (PDB: 8AHY). UCSF Chimera (*76*) was used for fitting of the separate rRic-8A-355/Nb8109/Nb8117/Nb8119 and NCS-1/Ric-8A-P in the cryo-EM map. Fitting of Ric-8A/Nbs was guided by the positioning of the Nbs, which acted as fiducials. Other docking poses yielded Nbs falling out of density (Fig S5B) Docking of NCS-1 was performed on the remaining density, testing all possible options (Fig S5B). The best fit was unambiguously assigned due to the best model filling all the available density as well as positioning the N-terminus of the Ric-8A-P pointing towards the connecting cryo-EM density. A light clash occurs between the Nb8119 and NCS-1 which is likely due to map inaccuracies at lower resolutions as well as conformational heterogeneity which cannot be modelled at the coordinate level with the current map resolution. Crude modelling of B8 and C8 into the unaccounted density was performed in Coot (*65*) using rigid body fitting.

### Guanine nucleotide exchange assays

Nucleotide exchange assays were carried out at 20 °C using a LS55 luminescence spectrometer (Perkin Elmer) with 5 nm slit widths (Ex/Em 295 nm/345 nm). Assays were conducted by measuring the change in rat ΔN31Gα_i1_ tryptophan fluorescence in the presence or absence of rRic-8A-491 or His-NCS-1/rRic-8A-491 as previously described (*39, 59*). All assays were conducted in 50 mM HEPES pH 8, 200 mM NaCl, 2 mM MgCl_2_, 1 mM TCEP. rRic-8A-491 or His-NCS-1/rRic-8A-491 were preincubated with rΔN31Gα_i1_ and 0, 1, 50 or 500 μM CaCl_2_ in a quartz fluorescent cuvette prior to addition of GTPγS (guanosine 5’-O-[gamma-thio]triphosphate. Final concentrations were as follows: 0.5 μM rRic-8A-491 or HisNCS-1/rRic-8A-491, 1 μM rΔN31Gα_i1_, and 10 μM GTPγS in 500 μl. For each assay a minimum of 6 technical repeats were performed. Progress curves were fit to a single or double exponential rate model and a one-way ANOVA followed by Dunnett’s multiple comparisons test was performed using GraphPad Prism (*77*). GTP binding rates of His-NCS-1/rRic-8A-491 were represented vs CaCl_2_ concentration. Data were fit to a One-site-total binding velocity model, *v = v*_*max*_ [Ca^2+^]/(K_a_ +[Ca^2+^]) + NS[Ca^2+^], where NS accounts for non-specific binding, using GraphPad Prism (*77*) to estimate an apparent Ca^2+^ activation constant (K_a_; mean ± SEM).

## Acknowledgements

M.J.S.-B. would like to thank ALBA (XALOC beamline) and ESRF synchrotrons for access and support of the staff, the mass spectrometry service from Institute “Rocasolano” and Prof. Armando Albert and Prof. Alberto Ferrús for critical revision of the manuscript. The proteomic analysis was performed in the proteomics facility of “Centro Nacional de Biotecnología” (CSIC). SEC-MALS experiments were performed at the Spectroscopy and NMR Unit (CNIO) with the assistance of Clara M. Santiveri and Ramón Campos-Olivas. This work was funded by grants from Spanish Ministry of Science and Innovation PID2019-111737RB-I00 (to M.J.S.-B.), PID2019-106608RB-I00 (to A.M.), RTI2018-099985-B-I00 (to M.M.) and PID2020-113359GA-I00 (to J.G.-N.). M.M. was supported also by CIBER of Respiratory Diseases (CIBERES) from ISCIII. A.M. and J.G.-N. were supported by “Ramón y Cajal” contracts from the Spanish Ministry of Science and Innovation (RYC-2017-22392 and RYC2018-025731-I, respectively). S.A.-U. was funded by a PhD fellowship of the “Diputación General de Aragón”. S.P.-S. was supported by a contract from “Programa de Empleo Juvenil de la Comunidad de Madrid” PEJ-2020-AI/BMD-18666. S.R.S. would like to acknowledge NIH grant P30GM140963 for support of the Center for Biomolecular Structure and Dynamics Integrated Structural Biology Core at the University of Montana, a Pilot Project grant to L.J.M., and R01GM105993 to S.R.S.

