## Supplemental Figures S1 to S7 and Movie S1 legend for "The neuronal calcium sensor NCS-1 regulates the phosphorylation state and activity of the Gα chaperone and GEF Ric-8A"

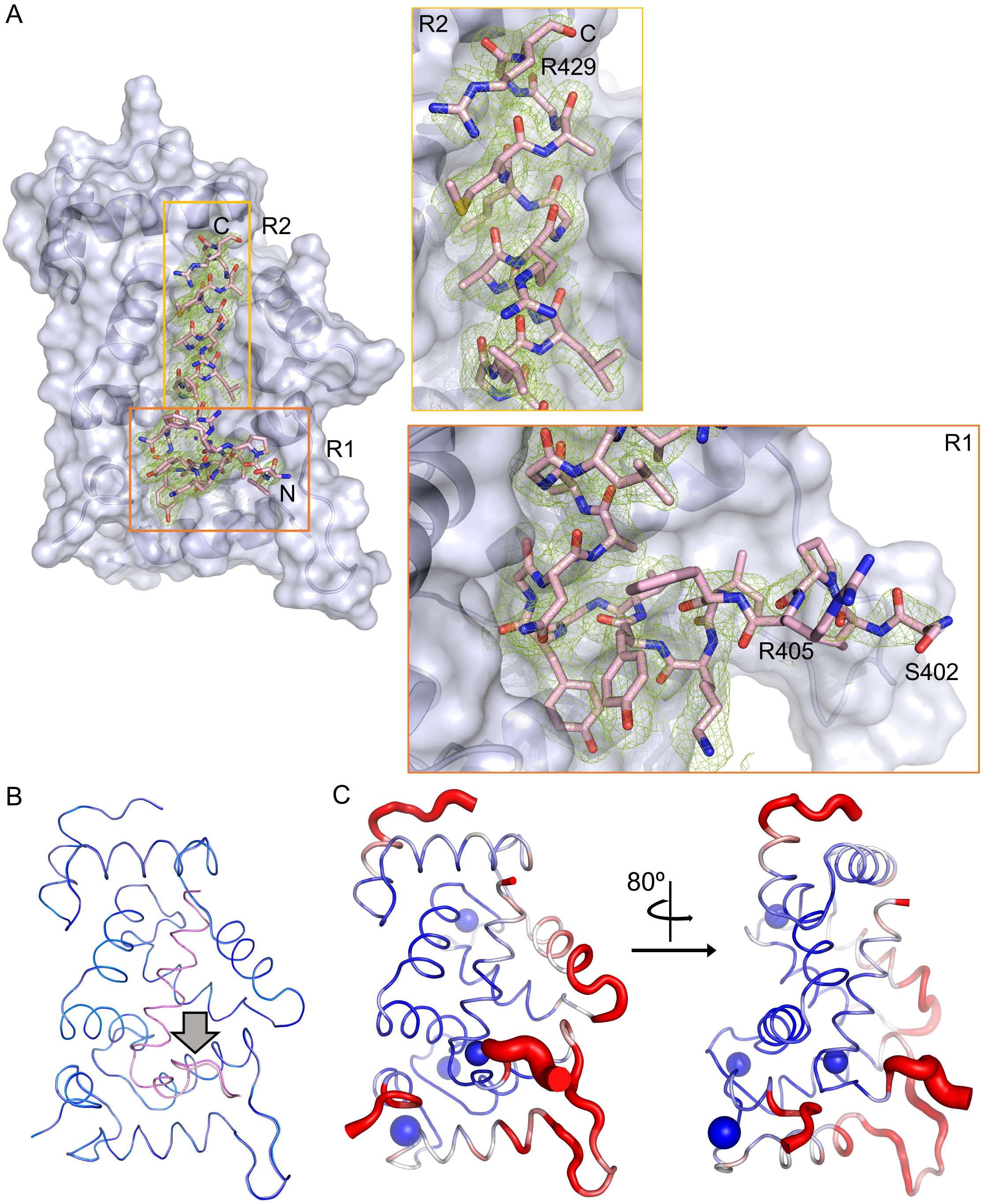


**Figure S1: Structure resolution of hRic-8A peptides bound to NCS-1**. **A)** Structure 1 showing the $2F_o$-$F_c$ electron density map (green) of Ric-8A-P2 (stick mode, pink). The molecular surface of NCS-1 is depicted. Squares represent magnifications of R1 and R2 regions. **B)** Left: Superimposition of Structure 1 and 2. NCS-1 and Ric-8A peptide threads in blue and in pink tones, respectively. The grey arrow indicates the main differences found in Ric-8A peptide structures. **C)** Temperature factor representation of Structure 2 (blue and red, high and low values, respectively) in two rotated views.

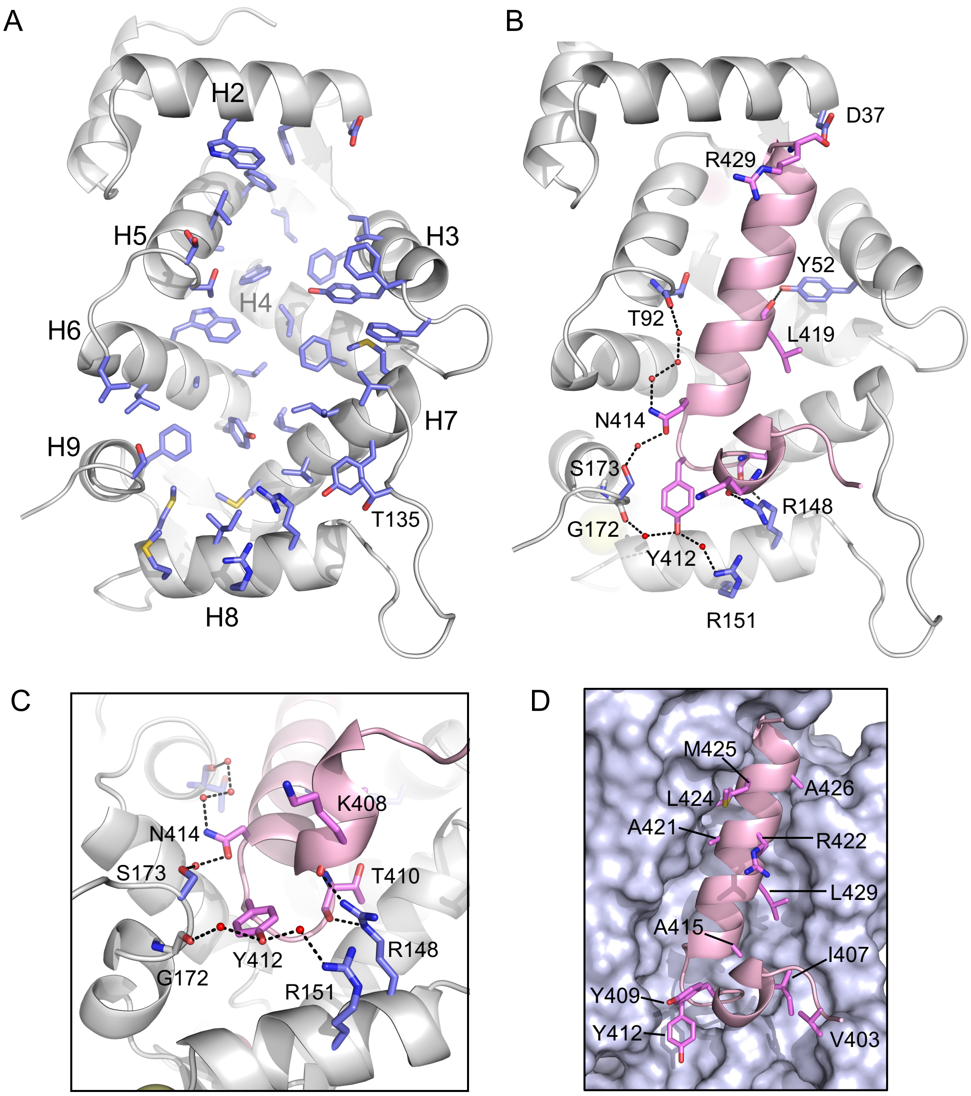


**Figure S2**: **The hNCS-1/Ric-8A-P protein-protein interface. A)** Ribbon representation of NCS-1. Helices are labelled and residues implicated in Ric-8A recognition are displayed as light purple sticks. **B)** H-bonds (black dashes) between NCS-1 (grey) and Ric-8A (pink). Interacting residues are shown as pink sticks and light-purple sticks, respectively. Water molecules are displayed as red spheres. **C)** H-bonds found in the R1-R2 loop. A rotated and zoomed view of that shown in B is depicted. **D)** Ric-8A residues implicated in van der Waals interactions are displayed in sticks and labelled. Ric-8A is shown as pink ribbon and the molecular surface of NCS-1 is represented.

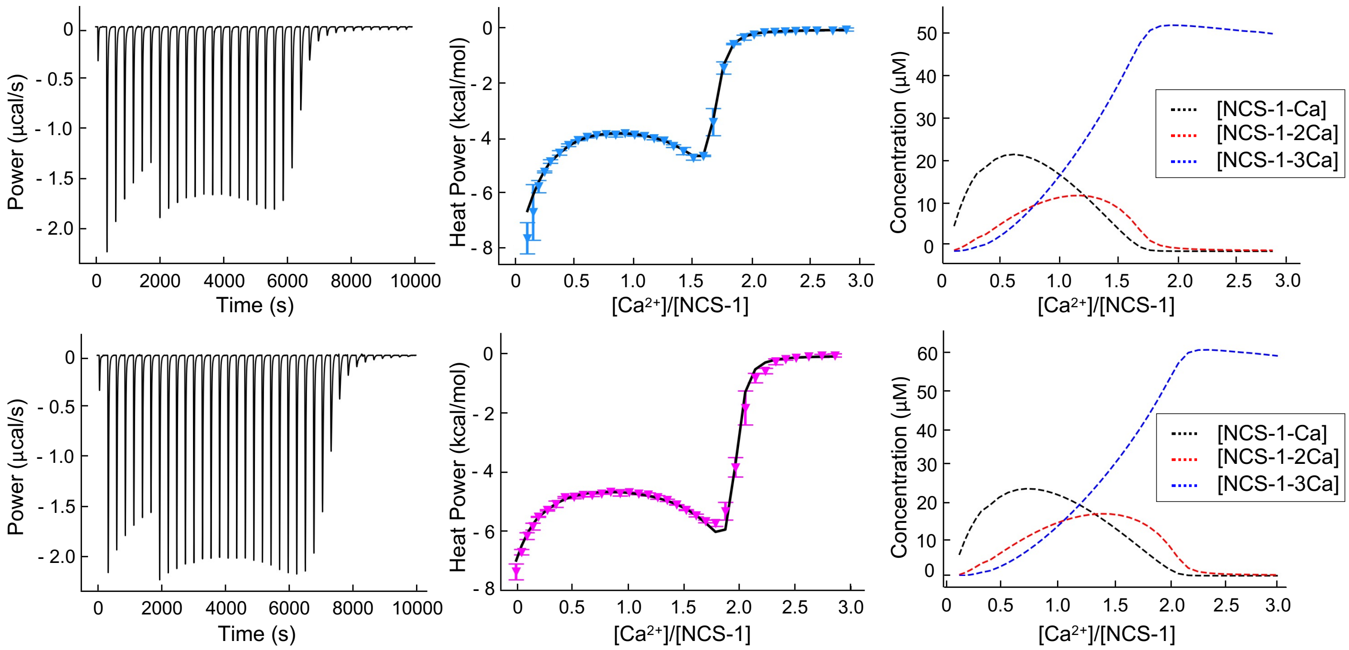


**Figure S3: Isothermal titration calorimetry.** Thermodynamics of $Ca^{2+}$ binding to NCS-1 in $Na^{+}$ (top panels) or $K^{+}$ (bottom panels) containing buffers. Experimental conditions as in Fig 4D. Raw data (left panels) and binding isotherms (central panels) in both conditions are shown, together with the NCS-1 population containing one, two, or three sites occupied with $Ca^{2+}$ as a function of the [Ca $^{2+}$]/[NCS-1] molar ratio (right panels). Best fits of titration isotherms using the three-site sequential binding model as shown as solid curves.

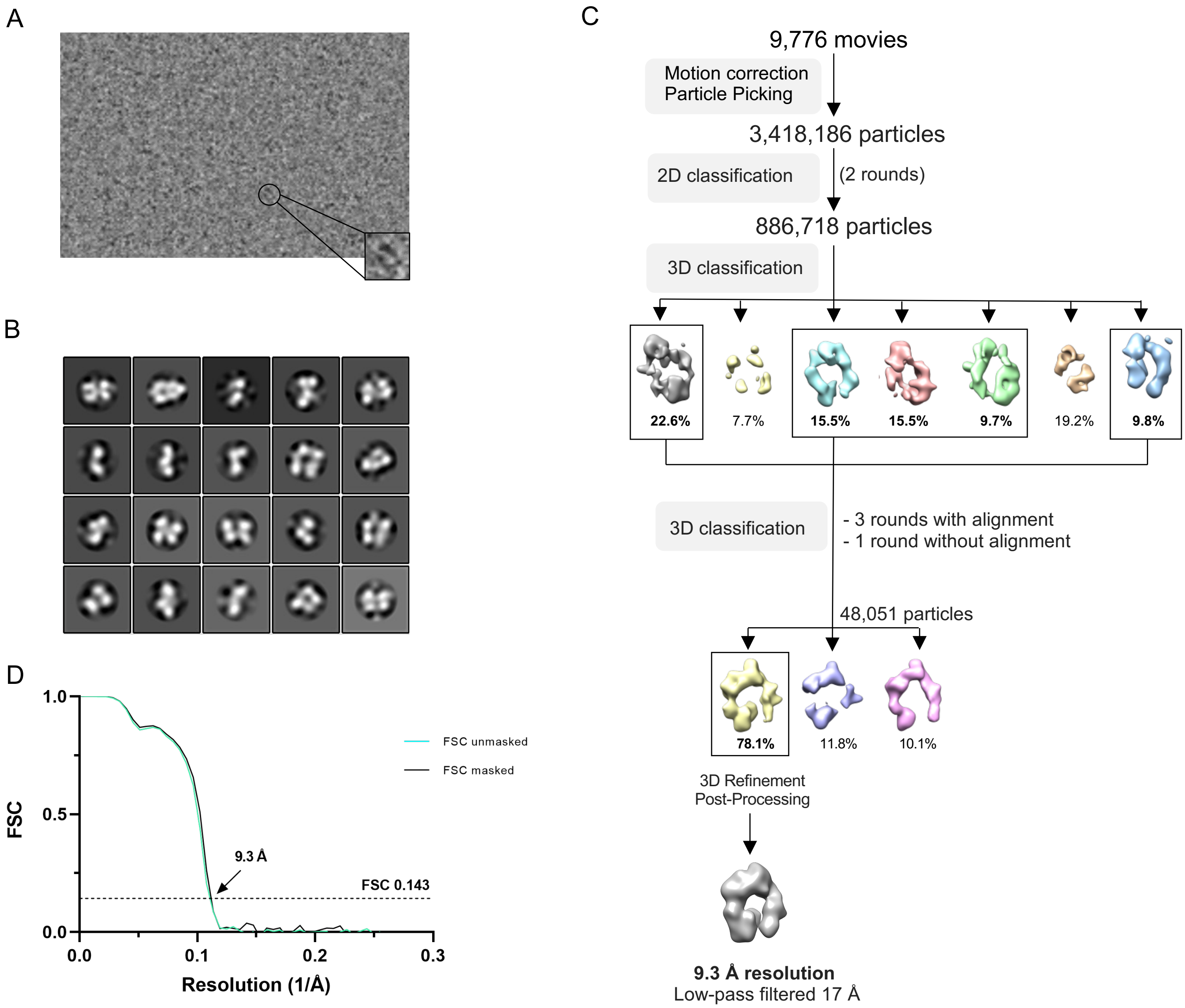


**Figure S4: Cryo-EM data processing workflow. A)** Representative Cryo-EM micrograph of NCS-1ΔH10/rRic-8A-452/Nb8109/Nb8117/Nb8119 at 0.6 mg/ml**. B)** 2D class averages of NCS-1ΔH10/rRic-8A-452/Nb8109/Nb8117/Nb8119. **C)** Data processing pipeline. **D)** FSC curves resulting from the refinement of two independent half-maps.

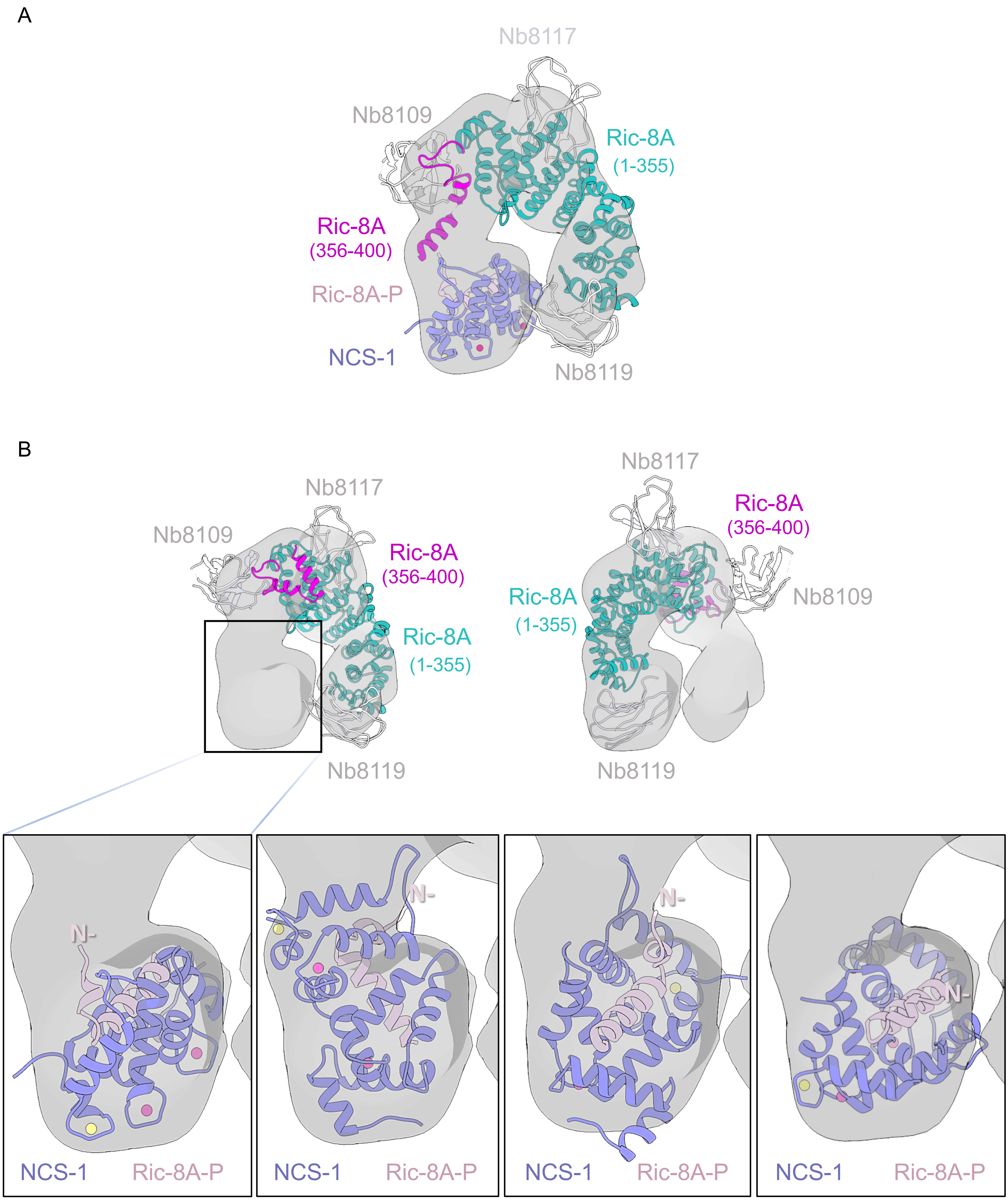


**Figure S5: Potential dockings of rRic-8A-400/Nbs and hNCS-1/Ric-8A-P complex in the cryo-EM map. A)** Hypothetical model of NCS-1/rRic-8A complex where A8-B8 loop and helices B8-C8 suffer a conformational change allowing this region to fit properly in the unaccounted density **B)** Front view showing a reasonable fit for rRic-8A-400/Nbs (PDB: 6UKT, (*39*)) (left panel) and an unlikely alternative docking of the complex (right panel). Inset to the remaining cryo-EM density with different hNCS-1/Ric-8A-P docking poses (lower panels). The left panel unveils the best fit with the N-terminal well oriented towards the extra-density. All structures are displayed as cartoons.

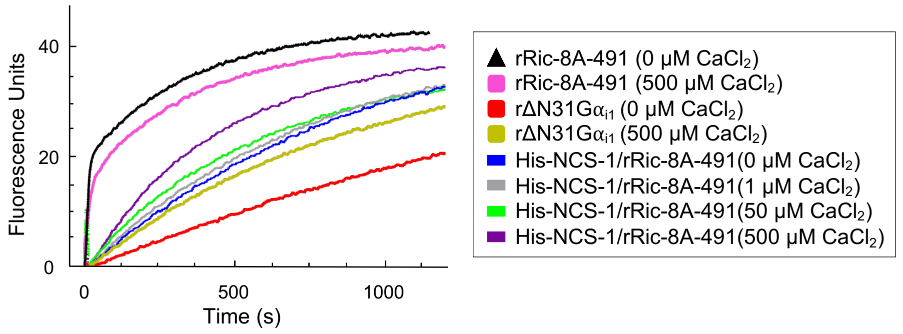


**Figure S6: GTPγS binding progress curves.** Representative traces of progress curves in determining nucleotide exchange rates shown in Fig 8. Change in fluorescence intensity is measured at excitation/emission = 295 nm/345 nm, which is sensitive to GTPγS binding $\Delta N31G\alpha_{i1}$.

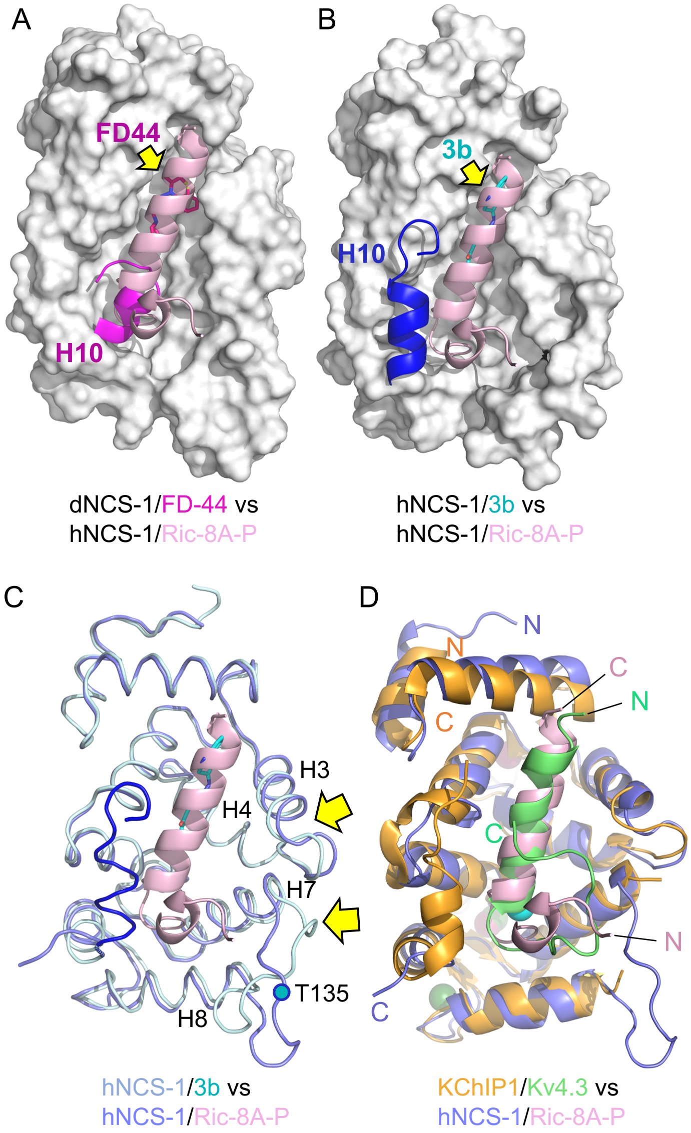


**Figure S7: Structural comparison of hNCS-1/Ric-8A-P with other NCS-1 complexes**. **A, B)** Superposition of NCS-1/Ric-8A-P (only Ric-8A is shown in pink ribbons) with other NCS-1 structures in complex with regulatory ligands, the PPI inhibitor FD-44 (*6*) and the PPI stabilizer 3b (*27*). The molecular surface of NCS-1 is represented except the helix H10 (ribbon). FD-44 and 3b compounds are represented in stick mode and yellow arrows indicate their position. **C)** Superposition of the structure of hNCS-1 (light purple) bound to Ric-8A-P (pink) with that of hNCS-1 (light blue; helix H10 in dark blue) bound to 3b regulator (cyan sticks). Yellow arrows indicate the NCS-1 regions that rearrange to accommodate Ric-8A. **D)** Superposition of the hNCS-1/Ric-8A-P (light purple/pink) complex with that of the KChIP1/Kv4.3 (orange/green) complex (PDB: 2I2R (*54*)). N- and C-terminal end of the different polypeptide chains are indicated following the same color code.

**Movie S1:** Morph movie explaining the structural rearrangement of Ric-8A HEAT repeat 9 for NCS-1 recognition. Ric-8A residues 402-429 are shown starting at the Gα-bound and ending at the NCS-1-bound conformations. The view is the same as that in Fig 7A. Side-chains are displayed in stick mode. While in the Ric-8A/Gα structure hydrophobic residues are at the back, facing the ARM-HEAT repeat domain (not shown), they rearrange and expose to the solvent to recognize NCS-1. The resulting structure is amphipathic and positively charged residues concentrate at the opposite side.
